# The legacy of maternal SARS-CoV-2 infection on the immunology of the neonate

**DOI:** 10.1101/2021.05.27.446014

**Authors:** Sarah Gee, Manju Chandiramani, Jeffrey Seow, Carlotta Modestini, Abhishek Das, Katie J Doores, Rachel M Tribe, Deena L Gibbons

## Abstract

Despite extensive and ongoing studies of SARS-CoV-2 and evidence that pregnant women are at increased risk of severe COVID-19, the effect of maternal infection on the developing infant remains unclear. To determine the potential impact of exposure to SARS-CoV-2 *in utero* on the neonate, we have assessed the immunological status of infants born to mothers with confirmed SARS-CoV-2 infection during gestation. No evidence of vertical transmission of SARS-CoV-2 was observed, but transfer of maternal SARS-CoV-2 specific IgG to infants was apparent, although to a lesser extent in cases of active or recent maternal infection. Infants born to mothers with recent/ongoing infection had elevated circulating pro-inflammatory cytokines and enhanced percentages of innate immune cells compared to that seen in infants born to uninfected mothers. In tandem, higher frequencies of FOXP3^+^ regulatory T cells and circulating IL-10 demonstrated a further nuance to the neonatal effector response. Interestingly, cytokine functionality was enhanced in infants born to mothers exposed to SARS-CoV-2 at any time during pregnancy. This indicates that maternal SARS-CoV-2 infection influences *in utero* priming of the fetal immune system.

## Introduction

Despite the ongoing COVID-19 pandemic, the effect of maternal SARS-CoV-2 infection on the immunology of the developing infant is still unclear. Indeed, SARS-CoV-2 infection in pregnancy has been reported to lead to variable outcomes for the mother. The majority of infected women in pregnancy are asymptomatic or only experience mild symptoms ^1 2^. Nevertheless, some pregnant women with SARS-CoV-2, especially in the third trimester, appear to be at an increased risk for hospitalization and subsequent intensive care unit admission ^3 4 5^ and rates of maternal infection increased in the second wave ^6 7^. Although the rates of preterm birth did not appear elevated in initial reports, more recent data suggest pregnant women are at a higher risk for subsequent preterm birth albeit much may be due to clinical intervention ^8^.

With respect to the infant, a national UK surveillance study suggested neonatal SARS-CoV-2 infection is uncommon even in babies born to mothers with perinatal infection ^9^. Similarly, a study from the US of 116 mothers with confirmed perinatal SARS-CoV-2 infection did not identify any neonatal cases ^10^ and a systematic review demonstrated no vertical transmission ^11^. Set against this, there have been a small number of individual case reports documenting evidence of vertical transmission^12 13 14 15 16,17^, although it was subsequently established that only 30% of documented cases were due to true placental transfer ^18^.

Whilst vertical transmission of SARS-CoV-2 itself is rare, the potential immunological perturbations induced in the pregnant mother ^19,20^ may conceivably leave an immunological legacy on the newborn with far reaching consequences. Indeed, our recent study in a preterm baby group found evidence of perinatal inflammation modulating the developing immune system of the infant ^21^. It is already appreciated that the immune system of the unborn child can be altered by the presence of human immunodeficiency virus (HIV) or Hepatitis C virus (HCV) in mothers, either with or without vertical transmission ^22 23 24 25^; that metabolites derived from maternal gut microbiota can shape the immune system of the offspring ^26^; and that modulation of the neonatal immune system has been associated with diseases in later life ^27 28^. In the limited studies that have assessed the immune status of babies born to SARS-CoV-2 infected mothers, there has been little evidence of impact in the cellular and humoral immunity of the neonate ^29^. Similarly, a small study of SARS-CoV-2 infection during pregnancy has been associated with a cytokine response in the fetal circulation (i.e. umbilical cord blood) but with no effect on the cellular immune repertoire ^20^. However, to our knowledge, none of the studies have included a comprehensive analysis of the infant cellular immune profile of babies born to SARS-CoV-2 exposed mothers (at any point in their pregnancy) compared to profiles of infants born to unexposed mothers. Indeed, it is increasingly challenging to include an appropriate control group in a global pandemic. Moreover, simultaneous analysis of cytokines and antibody titres in infants and their paired mothers is lacking. In our study, we have assessed the immunologic status of infants born to mothers with SARS-CoV-2 that tested positive either during the two weeks directly prior to birth or earlier on in pregnancy, compared to babies born to mothers never exposed to SARS-CoV-2 to identify if there is a legacy of maternal infection and potential *in utero* priming of the neonatal immune system.

## Results

### Transfer of passive immunity is reduced in infants born to mothers with recent/ongoing infection

Maternal and infant characteristics are shown in Table 1, and the group description is shown in Fig. 1a. Our neonatal group of infants born to mothers with SARS-CoV-2 exposure (SARS-CoV-2 exposed, SE, n=30) was divided into those with recent or ongoing infection as determined by mothers confirmed positive for SARS-CoV-2 by PCR within 2 weeks of birth (R/O, n=16, median positive swab 3 days before birth), or prior to 2 weeks; termed the recovered group (R, n=14, median positive swab 48.5 days prior to birth).

**Table 1.** Maternal and infant clinical characteristics.

| MATERNAL SARS-CoV-2 STATUS | SARS-CoV-2 Exposed |  | Non-SARS-CoV-2 Exposed (NSE) |
| --- | --- | --- | --- |
|  | Recent/Ongoing (R/O) | Recovered (R) |  |
| MATERNAL CHARACTERISTICS (n) | N=13 mothers | N=14 mothers | N=15 mothers |
| **Demographics** |  |  |  |
| <u>Age, median (range)</u> | 35 (26-43) | 32 (25-40) | 36 (30-46) |
| <u>BMI, median (range)</u> | 28 (18-42) | 28 (21-51) | 23 (18-30) |
| <u>Ethnicity</u> |  |  |  |
| White (British, Irish, Arab, European, African) | 10 | 9 | 12 |
| Black (African, Caribbean, British) | 5 | 4 | 2 |
| Asian | 0 | 1 | 0 |
| Mixed/Multiple Ethnicity | 0 | 0 | 1 |
| **COVID-related details** |  |  |  |
| <u>COVID symptoms, n</u> | 5 | 13 | N/A |
| <u>Cough</u> | 4 | 7 | N/A |
| <u>Fever</u> | 4 | 7 | N/A |
| <u>Shortness of breath (SOB)</u> | 3 | 6 | N/A |
| <u>Anosmia</u> | 1 | 2 | N/A |
| <u>Not documented</u> | 0 | 4 | N/A |
| <u>Asymptomatic</u> | 10 | 1 | N/A |
| <u>Days between positive SARS-CoV-2<sup>+</sup> status (nasopharyngeal swab) and birth, median (range)</u> | 3 (0-13) | 48.5 (22-221) | N/A |
| <u>Gestation (weeks<sup>+days</sup>) at time of positive SARS-CoV-2<sup>+</sup> status, median (range)</u> | 37<sup>+6</sup> (31<sup>+4</sup>-41<sup>+5</sup>) | 33<sup>+1</sup> (7<sup>+0</sup>-36<sup>+4</sup>) | N/A |
| **Comorbidities (n)** |  |  |  |
| <u>Diabetes</u> |  |  |  |
| Gestational diabetes mellitus (GDM) | 2 | 3 | 1 |
| Type 2 diabetes mellitus (T2DM) | 1 | 0 | 0 |
| No diabetes | 12 | 11 | 14 |
| <u>Hypertension</u> |  |  |  |
| Chronic | 1 | 0 | 0 |
| Gestational/current | 0 | 1 | 1 |
| Postnatal | 1 | 0 | 0 |
| Unknown | 0 | 0 | 1 |
| None | 13 | 13 | 13 |
| <u>Asthma</u> | 0 | 1 | 3 |
| <u>Co-infection</u> |  |  |  |
| Group B streptococcus (GBS) | 1 | 3 | 0 |
| Primary HSV | 1 | 0 | 0 |
| None | 13 | 11 | 15 |
| <u>Acute Chorioamnionitis/Subchorionitis</u> | 1 | 3 | 0 |
| <u>Ankylosing spondylitis</u> | 0 | 1 | 0 |
| **Delivery information** |  |  |  |
| <u>Pyrexia</u> | 1 | 1 | 0 |
| <u>No pyrexia</u> | 14 | 13 | 15 |
| **INFANT CHARACTERISTICS (n)** | N=16 infants | N=14 infants | N=15 infants |
| <u>Singletons (n)</u> | 14 | 14 | 15 |
| <u>Twins (n)</u> | 2 (1 set) | 0 | 0 |
| <u>Mode of Delivery (MOD)</u> |  |  |  |
| Caesarean Section (CS) | 10 | 8 | 15 |
| Vaginal Delivery (VD) | 6 | 6 | 0 |
| <u>Gestational Age</u> |  |  |  |
| >37 weeks | 11 | 13 | 15 |
| 36-37 weeks | 3 | 1 |  |
| <34 weeks | 2 | 0 | 0 |
| <u>Sex</u> |  |  |  |
| Male | 9 | 8 | 8 |
| Female | 7 | 6 | 7 |
| <u>Apgar Scores at 5 mins, median (range)</u> | 9 (8-10) | 10 (9-10) | Not known |

**Fig. 1.**
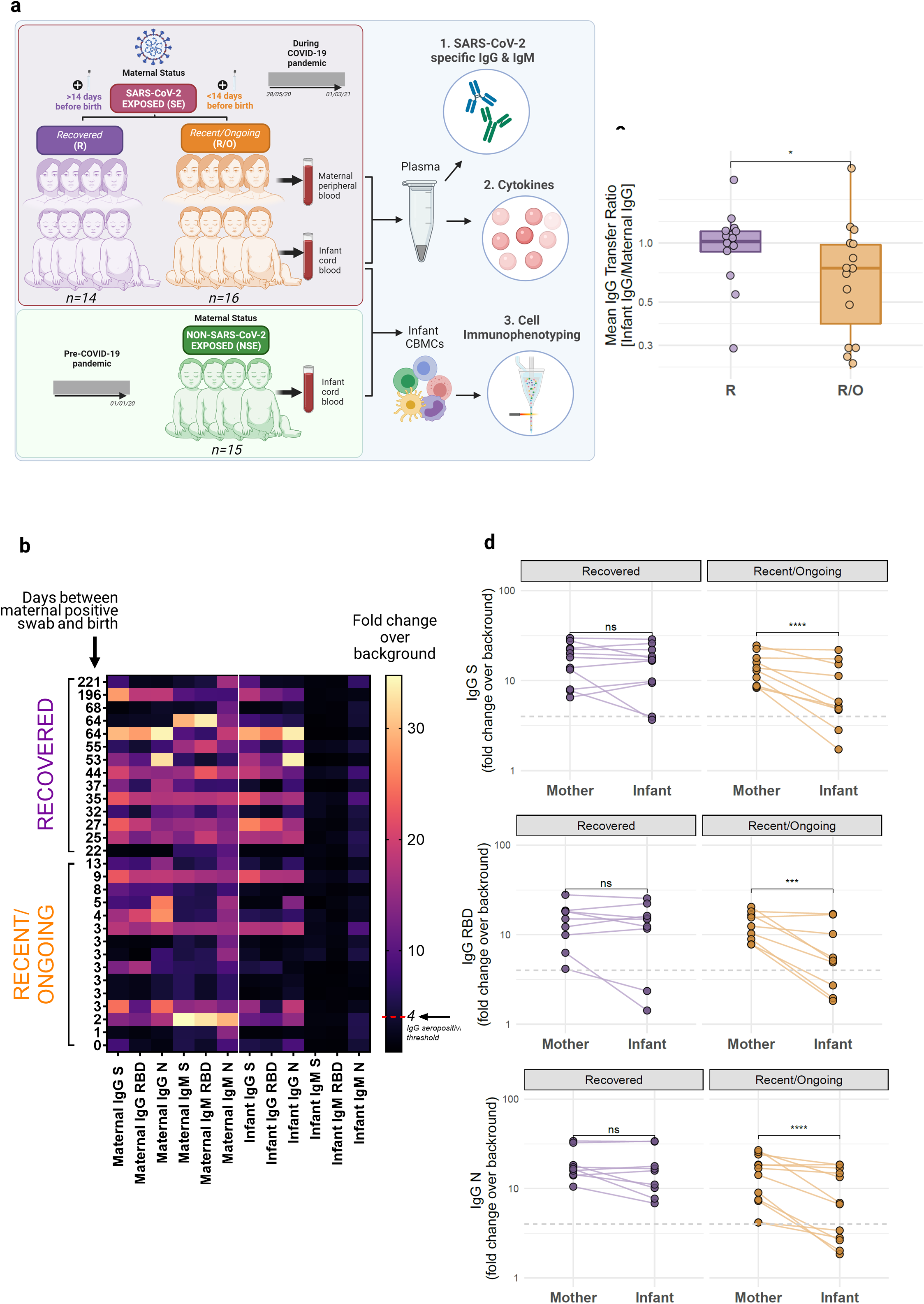
Infants born to SARS-CoV-2 exposed mothers are IgG^+^IgM^-^ and reduced antibody transfer is seen in the recent/ongoing group. **a**, Study outline illustrating recruitment of infants and mothers in the SE group (*n=30*), comprised of two groups (R: *n=14*, R/O: *n=16*), and infants in the NSE group (*n=15*). Figure created with BioRender.com. **b**, Heatmap displaying paired maternal and infant IgG and IgM levels against S, RBD and N SARS-CoV-2 epitopes measured in plasma from maternal blood and cord blood taken at the time of delivery. Peak IgG/IgM levels (fold change over background) are displayed within the infant-mother dyads (each row, R: *n=14*, R/O: *n=15*), ordered by the number of days between maternal positive nasopharyngeal swab and birth. 4 fold change (over background) is defined as the IgG seropositivity threshold. **c**, Boxplot of the mean IgG (average of IgG S, N and RBD) transfer ratio between the infants and their paired mothers within the R group (*n=14*) and R/O group (*n=15*). **d**, Maternal and infant paired peak IgG levels (fold change over background) in dyads with seropositive mothers only (4 fold change over background = dashed grey line) within the R (IgG S: *n=11*, IgG RBD: *n=9*, IgG N: *n=9*) and R/O (IgG S: *n=10*, IgG RBD: *n=9*, IgG N: *n=12*) groups. Each line joins an infant and their paired mother. Boxplots follow standard Tukey representations; central line = median, upper line = 75^th^ percentile; lower line = 25^th^ line; whiskers = 1.5 x 75^th^/25^th^ percentile. *p<0.05; **p<0.01; ****p<0.0001 assessed by two-sided Wilcoxon rank-sum tests (**c**) and two-sided paired (**d**) Wilcoxon tests.

In both the recent/ongoing and recovered group, regardless of the time between the positive swab and birth or maternal IgM levels, SARS-CoV-2 specific IgM was not detected in cord plasma, suggestive of a lack of vertical transmission (Fig. 1b). This was true for IgM directed against the spike protein (S), the receptor binding domain within the spike protein (RBD) as well as the nucleoprotein (N) for which background IgM reactivity has been shown to be higher ^30^. By contrast, SARS-CoV-2 specific IgG against all three antigens was detected in infants born to SARS-CoV-2 exposed mothers (Fig. 1b). Whilst levels of SARS-CoV-2 specific IgG in the mother-infant dyad were comparable in the recovered group, there were significantly lower levels of SARS-CoV-2 specific IgG in infants born to mothers with recent/ongoing infection (Extended Data Fig. 1a). Thus, when the ratio of infant Ig to their paired maternal Ig (transfer ratio) was calculated for each antigen, the mean transfer ratio of all 3 antigens was 1.04 in the R group and 0.79 in the R/O group (Fig. 1c). This was despite the presence of high levels of maternal IgG in at least some mothers in the R/O group (Extended Data Fig. 1a). Indeed, when only comparing IgG levels in infants born to seropositive mothers, transfer of SARS-CoV-2 specific IgG to the infant was still significantly lower in the R/O group (Fig. 1d) and this did not appear to differ with the sex of the infant (Extended Data Fig. 1b).

### Elevated plasma cytokines in mothers with recent/ongoing infection and their infants

SARS-CoV-2 infection is known to be associated with marked elevation of several plasma cytokines including Interferon gamma-induced protein 10 (IP-10), interleukin (IL)-1β, CXCL8, IL-6 and IL-10 ^31,32 33 34^. To assess the impact of this on the neonate, plasma cytokine concentrations were assessed in paired maternal and cord blood using a multiplex assay. IP-10 and IL-1β levels in the plasma of mothers with recent/ongoing SARS-CoV-2 infection were significantly elevated when compared to that of recovered mothers whilst IL-10, CXCL8 and IL-6 were similar between the SE groups (Fig. 2a). When assessing neonatal cytokine levels, IL-10 was significantly elevated in the cord plasma from babies born to mothers with recent/ongoing infection compared to those born to recovered mothers. CXCL8 levels were also numerically higher in the recent/ongoing group although this did not reach significance, largely driven by three infants with undetectably levels (Fig. 2b). However, concentrations of this chemokine was significantly higher in infants than in their paired mothers (Fig. 2c) which was not seen with any of the other cytokines tested. Conversely, infant IP-10 was significantly lower than their paired mothers (Extended Data Fig. 2a). The majority of babies born to recovered mothers that showed elevated levels of CXCL8 were born by vaginal delivery (Fig. 2d), known to elevate several cytokines ^35^. However, there were still notable increases of CXCL8 in babies born via caesarean section (CS) in the recent/ongoing group, compared to the recovered group, suggesting this was indeed a fetal response to maternal SARS-CoV-2 infection.

**Fig. 2.**
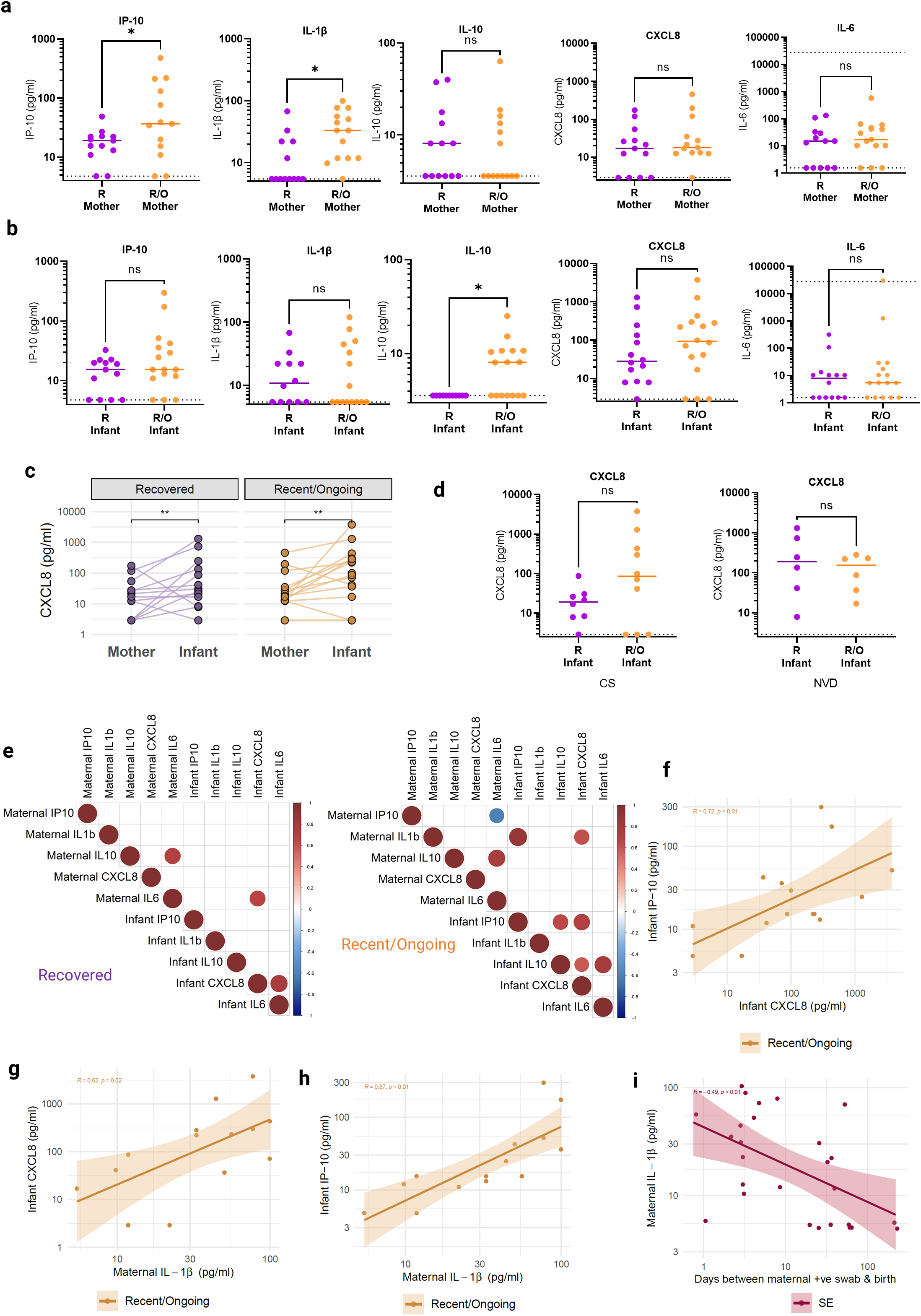
Elevated cytokine levels in plasma from infant cord blood and paired maternal peripheral blood within the recent/ongoing group. **a-b**, IP-10, IL-1β, IL-10, CXCL8 and IL-6 levels in the mothers and their infants (**b**) within the R (Mothers - *n=13*, Infants - CXCL8 & IL-6: *n=14*, IP-10: *n=13*, IL-1β: *n=12*, IL-10: *n=11*) and the R/O (Mothers – IL-6: *n=15*, IL-10 & IL-1β: *n=14*, CXCL8 & IP-10: *n=13*, Infants – IP-10, CXCL8, IL-10, IL-6: *n=16*, IL-1b: *n=15*) groups measured in plasma from maternal blood and cord blood taken at the time of delivery. **c**, Paired maternal and infant plasma CXCL8 levels within the R and the R/O groups. Each line joins an infant and their paired mother. Horizontal dotted lines represent the minimum (and maximum, for IL-6) detectable concentrations. **d**, Spearman correlation matrices of all significant (p<0.05) correlations of infant and maternal cytokines in the R (LHS: *n=14*) and R/O (RHS: *n=16*) groups. **e-h**, Spearman correlation plots with generalised linear model lines and 95% confidence intervals in the R/O group (**e-g**) (e: *n=15*, f-g: *n=14*) and the SE group (**h**) (*n=27*). **i**, Infant CXCL8 levels in babies born via CS (LHS: *n=18*) or NVD (RHS: *n=12*). Cytokine scatter plots central line = median value. Unadjusted p values (*p<0.05; **p<0.01) were assessed by Kolmogorov-Smirnov tests (**a**,**b**,**i**) and two-sided paired Wilcoxon tests (**c**).

Spearman rank test identified significant (p<0.05) correlations in cytokine levels both within and between maternal and cord blood in both groups. Nonetheless, this was more evident in mothers and infants in the R/O group compared with the recovered group suggestive of a greater degree of immune co-regulation in recent/ongoing SARS-CoV-2 infection (Fig. 2e): there was a correlation between CXCL8 and IP-10 in infants born to mothers with recent/ongoing infection (Fig. 2f); maternal IL-1β levels correlated with CXCL8 and IP-10 in the infants from this group (Fig. 2g-h), and interestingly, maternal IL-1β levels negatively correlated with days between a positive COVID swab and birth (Fig. 2i), suggesting this cytokine was indicative of recent infection in the mothers. No differences were seen in plasma IL-12p70, GM-CSF, IFN-α2, IFN-λ1, IFN-λ2/3, IFN-β, TNF-α and IFN-γ in mothers or infants from either group (Extended Data Fig. 2b).

### Recent/ongoing maternal SARS-CoV-2 infection influences adaptive immune cell populations in the infant

To determine the potential impact of maternal SARS-CoV-2 infection on the cellular immune compartment of the neonate at birth, we employed multiparametric flow cytometry to phenotype peripheral blood leukocytes and assess their *in vitro* functional capacity upon mitogen stimulation (Figs. 3-5). Gating strategies are shown in Extended Data Fig. 3. To establish if maternal SARS-CoV-2 infection altered the developing immune system of the neonate, infant cellular immune profiles in the combined (R and R/O) SARS-CoV-2 exposed group (SE; n=30) were compared to those from term infants born to healthy mothers collected prior to the pandemic (Non SARS-CoV-2 exposed, NSE; n=15) but measured contemporaneously. We performed tSNE dimensionality reduction (Fig. 3a) on 91 individual flow cytometry immune parameters and observed that infant immune profiles in the SE group clustered away from the normal immune profiles of the NSE group. Correlations between immune parameters appeared significantly different in infants born to SE mothers compared to NSE mothers (Fig. 3b-c). For example, a weaker correlation was seen between IFN-γ producing cells and a stronger correlation occurred between proliferating TEMRAs (Extended Data Fig. 4a-b). Indeed, when focusing on the immune parameters that drive the biggest differences, we identified that infants born to mothers in the R/O group separated furthest away from the NSE group, using 3-dimensonal PCA (Fig. 3d), and that all three groups tended to segregate based on their maternal SARS-CoV-2 status upon unbiased hierarchal clustering analysis (Fig. 3e). Moreover, there was no clear clustering based on alternative confounding factors such as sex, ethnicity, mode of delivery and other maternal characteristics such as chorioamnionitis and gestational diabetes (GDM) (Extended Data Fig. 5). Despite the segregation observed in the infant cellular immune compartment, many parameters, including those known to be perturbed in the adult response to SARS-CoV-2, were not different between groups. Infant T cell lymphopenia was not observed and moreover the relative frequencies of major adaptive lymphocyte subsets (e.g. CD4 and CD8 αβ T cells, γδ T cells and B cells) were unaffected by maternal exposure to SARS-CoV-2 (Fig. 3f). Despite preserved composition of cell frequencies, changes in adaptive cell populations were still evident, including an increase in the percentage of CD161 expressing CD8 T cells in babies born to mothers with recent/ongoing infection (Fig. 3g) as well as increased CD25^+^FOXP3^+^ T_REGS_ (Fig. 3h), the latter of which positively correlated with the percentage of Vδ2 T cells (Fig. 3i).

**Fig. 3.**
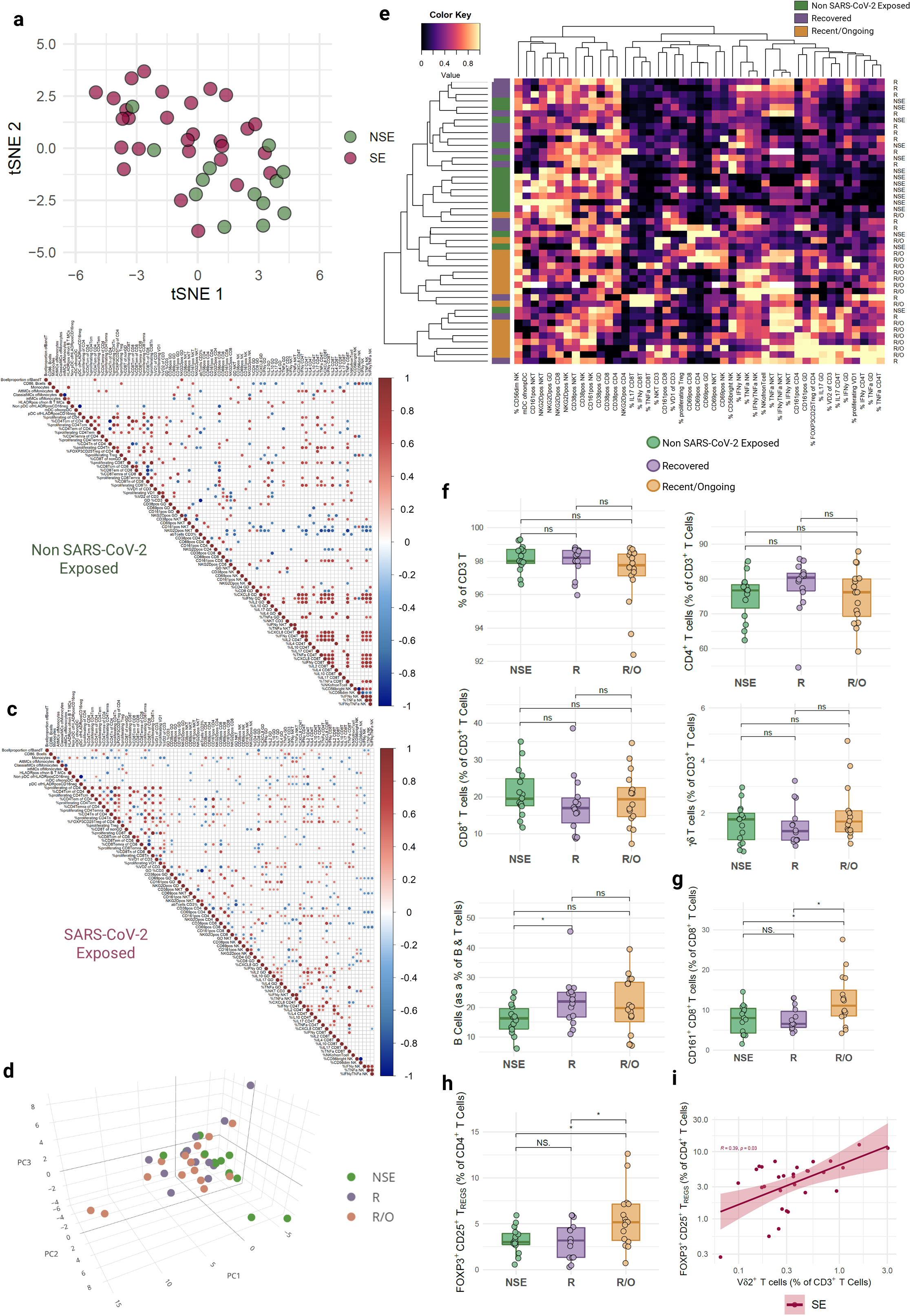
Recent/ongoing maternal SARS-CoV-2 infection influences some, but not all, adaptive immune cell proportions in infant cord blood. **a**, tSNE dimensionality reduction of infant immune profiles measured from cord blood mononuclear cells (CBMCs) in those born to SARS-CoV-2 Exposed mothers (SE; *n=28*) and Non SARS-CoV-2 Exposed mothers (NSE; *n=13*) infants. **b**,**c**, Spearman correlation matrix of significant (p<0.05) correlations of infant cellular populations in the NSE (*n=15*) the SE (*n=30*) (**c**) groups. **d**, 3D PCA dimensionality reduction of infant immune profiles in the Recent/Ongoing group (R/O; *n=15*), Recovered group (R; *n=13*) and Non SARS-CoV-2 Exposed infant groups (NSE; *n=13)*. PC1, PC2, and PC3 explain 20.6%, 11.8% and 8.4% of the variance, respectively. **e**, Dendrogram cluster heatmap of 45 flow cytometry immune populations in infants within the NSE (*n=14*), R (*n=14*), R/O (*n=15*) groups. **f**, Boxplots displaying the proportions of αβ T cells, CD4^+^ T cells, CD8 T cells and γδ T cells in the (NSE: *n=15*, R: *n=14*, R/O: *n=16*) groups, and proportion of B cells (NSE: *n=14*, R: *n=14*, R/O: *n=16*). **g-h**, Boxplots displaying the proportions of CD161^+^CD8^+^ T cells and FOXP3^+^CD25^+^ T_REG_ cells (NSE: *n=15*, R: *n=14*, R/O: *n=16*). **i**, Spearman correlation plots in the SE group (*n=30*) of infant proportions of FOXP3^+^CD25^+^ T_REG_ cells and Vδ2 γδ T cells with generalised linear model lines and 95% confidence intervals. All boxplots follow standard Tukey representations; central line = median, upper line = 75^th^ percentile; lower line = 25^th^ line; whiskers = 1.5*75^th^/25^th^ percentile. Unadjusted p values (*p<0.05) were assessed by two-sided Wilcoxon rank-sum tests.

**Fig. 4.**
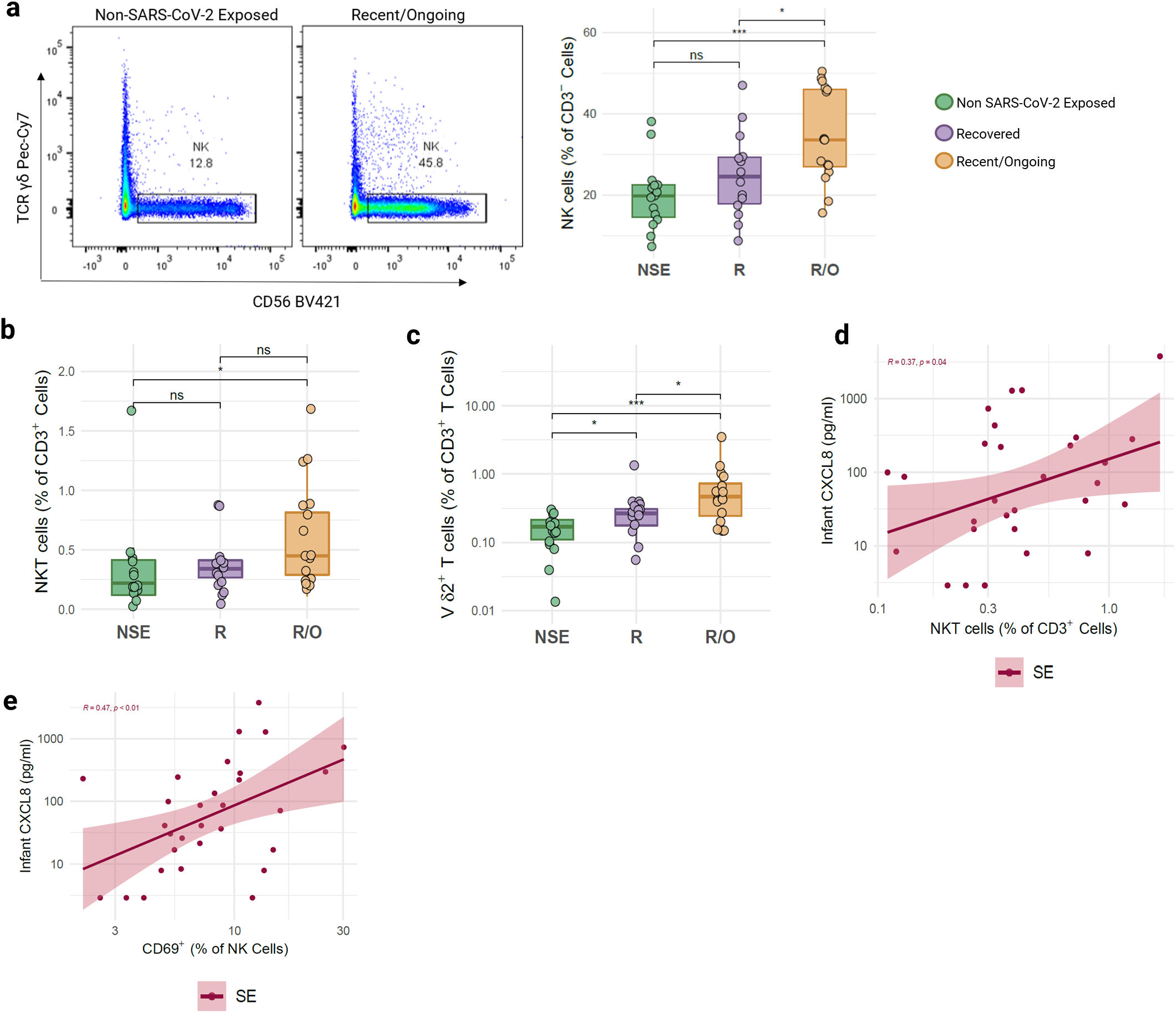
Elevated proportions of innate immune cells in cord blood from infants born to mothers with recent/ongoing SARS-CoV-2 infection. a, Representative flow cytometry plots of % CD56^+^ NK cells (of CD3^-^ live cells) in an NSE and R/O infant, measured from cord blood mononuclear cells (CBMCs), and the boxplot for the total infants (NSE: *n=15*), R: *n=14*), R/O: *n=16*). **b-c**, Boxplots displaying the proportions of NKT cells and Vδ2 γδ T cells (NSE: *n=15*, R: *n=14*, R/O: *n=16*). **d-e**, Spearman correlation plots in the SE group t (*n=30*) of infant CXCL8 levels and the proportions of NKT cells (**d**) or CD69^+^ NK cells (**e**) with generalised linear model lines and 95% confidence intervals. All boxplots follow standard Tukey representations; central line = median, upper line = 75^th^ percentile; lower line = 25^th^ line; whiskers = 1.5*75^th^/25^th^ percentile. Unadjusted p values (*p<0.05; ***P<0.001) were assessed by two-sided Wilcoxon rank-sum tests.

**Fig. 5.**
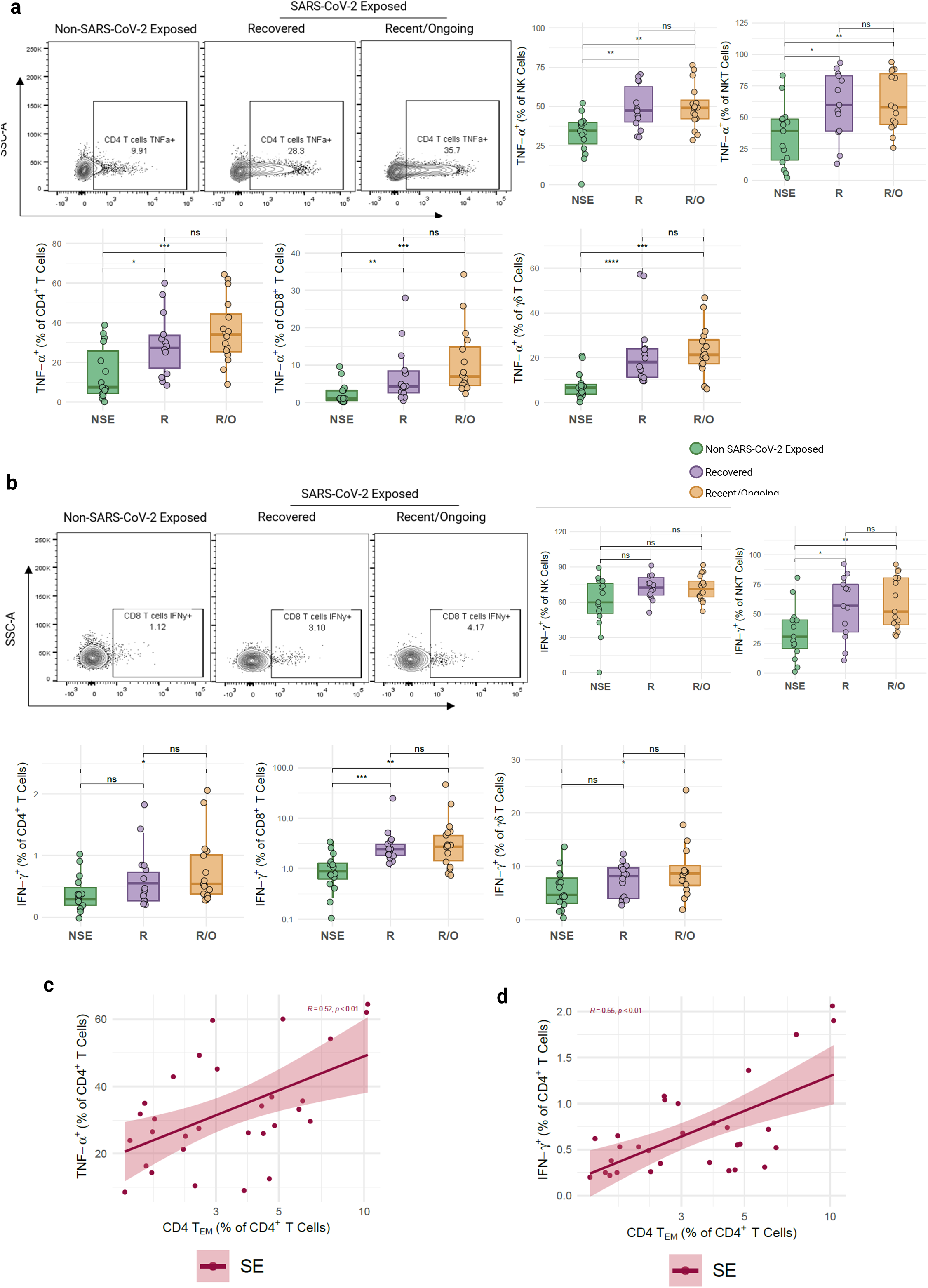
Increased cell cytokine functionality following polyclonal stimulation of CBMCs from infants born to SARS-CoV-2 exposed mothers. **a**,**b**, Representative flow cytometry plots of % TNF-α^+^ CD4^+^ T cells (**a**) and % IFN-γ^+^ CD8^+^ T cells (**b**) in an NSE, R and R/O infant, measured from cord blood mononuclear cells (CBMCs). Boxplots displaying the proportions of TNF-α^+^ (**a**) and IFN-γ^+^ (**b**) CD4^+^ T cells, CD8^+^ T cells, γδ T cells and NK cells (NSE: *n=15*, R: *n=14*, R/O: *n=16*), as well as NKT cells (NSE: *n=15*, R: *n=13*, R/O: *n=15*), following polyclonal stimulation [PMA (10 ng/ml), Ionomycin (1 μg/ml), Brefeldin A (20 ng/ml) and Monensin solution (2 μM) at 37°C for 4 h]. **c-d**, Spearman correlation plots of infant CD4^+^ T_EM_ cells/TNF-α^+^CD4^+^T (**c**), CD4^+^ T_EM_ cells /IFN-γ^+^CD4^+^T cells (**d**) in the SE group (*n=30*) with generalised linear model lines and 95% confidence intervals. All boxplots follow standard Tukey representations; central line = median, upper line = 75^th^ percentile; lower line = 25^th^ line; whiskers = 1.5*75^th^/25^th^ percentile. Unadjusted p values (*p<0.05; **p<0.01; ***P<0.001) were assessed by two-sided Wilcoxon rank-sum tests.

### Conspicuous increased proportions of innate immune cells in neonates born to mothers with recent/ongoing SARS-CoV-2 infection

In contrast to the adaptive immune cell compartment, changes in innate cell subsets were more prominent. NK, NKT and innate-like Vδ2 γδ T cells were significantly elevated in babies born to mothers with recent/ongoing infection (Fig. 4a-c). There was also a change in monocyte populations with enhanced percentages of alternative monocytes and subsequently reduced CD38^+^ classical monocytes in babies born to mothers with recent/ongoing infection (Extended Data Fig. 6a). These cells were, for the most part, not significantly elevated in infants born to mothers with previous SARS-CoV-2 exposure (R), consistent with reactive neonatal responses to recent/ongoing SARS-CoV-2 infection. Indeed, raised percentages of NK cells negatively correlated with days from positive SARS-CoV-2 swab result to birth, further suggesting this was a neonatal response to maternal infection (Extended Data Fig. 6b). Interestingly, the percentage of NKT cells (Fig. 4d) and NK cell activation (as assessed by CD69 expression, Fig. 4e) both positively correlated with the levels of cord blood CXCL8 suggesting these are key immune markers associated with the neonatal response to maternal infection.

### Increased cytokine functionality in innate and adaptive cells in all infants born to SARS-CoV-2 exposed mothers

When assessing functionality by intracellular cytokine staining post polyclonal activation, the ability of immune cells to produce cytokines upon stimulation was significantly elevated in babies born to SE mothers. Consistent with changes observed in plasma cytokine concentrations and cellular immune composition, enhanced cytokine production by neonatal immune cells was associated with maternal SARS-CoV-2 infection. Indeed, cytokine potential was significantly enhanced in several different cell types in babies born to mothers previously exposed to SARS-CoV-2 (at any time point) as exemplified by TNF-α (Fig 5a), IFN-γ (Fig. 5b) and to a lesser extent, IL-17 (Extended Data Fig. 6c) although the ability to produce CXCL8 was unaltered (Extended Data Fig. 6d). The observed enhanced cytokine functionality (TNF-α and IFN-γ) in CD4 T cells positively correlated with effector memory CD4 T cells (Fig. 5 c,d), and TNF-α producing cells (both CD4 and CD8) negatively correlated with CD38 expression, known to decrease during maturation (Extended Data Fig. 6e) ^21^. Due to limited numbers of mothers with severe disease, we were unable to establish if the extent of immune imprinting was related to maternal infection status.

## Discussion

Our study provides a comprehensive immune atlas of neonates born to mothers with SARS-CoV-2 exposure. Whilst we did not observe vertical transmission of SARS-CoV-2 itself, we did find multiple immunological perturbations within the neonate associated with maternal SARS-CoV-2 exposure during pregnancy, many of which were also associated with recent/ongoing infections. Taken together, our findings are suggestive of an immunological legacy imprinted on the neonate following maternal SARS-CoV-2 exposure, with potential far-reaching consequences. *In utero* exposure to environmental factors, infection and/or maternal inflammation is increasingly recognised to affect the developing immune system and subsequent responses both to infection ^36^, immune mediated diseases ^37-39^ and neurodevelopmental problems ^40,41^. Indeed, long term effects cannot be ruled out as observed in survivors after *in utero* exposure to the 1918 (Spanish) influenza pandemic ^42 43^.

Although we did not directly assess neonates for the presence of SARS-CoV-2, we did assess SARS-CoV-2 specific IgM levels and could find no evidence of vertical transmission in any of the 30 infants born to SARS-CoV-2 exposed mothers. SARS-CoV-2 specific IgG was, however, transferred to the neonates from their mothers suggestive of the transfer of protective immunity. There was a correlation between maternal and infant SARS-CoV-2 IgG levels in mother-infant dyads in both groups as previously suggested ^44^. However, even though for many pathogens, umbilical cord titers of IgG at normal term delivery are higher than in maternal blood ^45 46 47 48^, there were reduced levels of SARS-CoV-2 specific IgG in infants born to mothers with recent/ongoing infection compared to their paired mothers. This did not appear to be a threshold issue, as many mothers exhibited high levels of SARS-CoV-2 specific IgG which was not transferred efficiently to their infant. Reduction of SARS-CoV-2 specific Ig transfer via the placenta has been suggested to occur in the third trimester due to altered glycosylation ^49^ and reduced maternal SARS-CoV-2-specific antibody titers and impaired placental antibody transfer were also noted in pregnancies with a male fetus ^50^, although there did not appear to be any sex bias in this data set. It is currently unclear whether antibodies induced via vaccination as opposed to natural infection, differ in terms of their glycosylation status and subsequent placental transfer. Vaccination to SARS-CoV-2 in the second and third trimester did elicit placental transfer of Abs, with a reduced transfer ratio observed in the last trimester^51 52^.. Our study adds further evidence suggesting the 2^nd^ trimester may represent more opportune vaccination timing, at least with respect to the transfer of passive immunity to the infant.

Perhaps unsurprisingly, cord plasma of neonates born to mothers with recent/ongoing infection, expressed elevated concentrations of some cytokines known to be associated with inflammation and COVID-19, consistent with placental immune activation ^31^. Previously, elevated levels of IP-10, IL-6, IL-10, CXCL8 and IL-1β have been associated with adult infection and the former three with disease severity ^32 33 34^. Increased IL-6 and IL-10 have also been associated with severity in early SARS-CoV-2 infection in children ^53^. Interestingly, this conventional COVID-19 signature of adults was skewed more towards IL-10 (and to a lesser extent, CXCL8) in the neonates born to mothers with recent/ongoing infection. Theoretically, the increased concentrations of cytokines in the infant could be explained by transfer of maternal cytokines through the placental tissues. Indeed, infants with raised IP-10 concentrations were born to mothers who had high plasma IP-10 measurements, although in general concentrations were significantly lower in infants compared to mothers. However, in the case of cord plasma CXCL8, a chemokine which correlated strongly with maternal IL-1β (previously associated with maternal SARS-CoV-2 infection, ^54^) levels were significantly higher than that observed in their mothers, suggesting that at least some of these elevated cytokines were fetal-derived, and a direct response to maternal infection. Elevated cord plasma cytokines, including CXCL8, have been observed in some infants born to SARS-CoV-2^+^ mothers ^20^. Similarly, increased cord plasma CXCL8 has been observed after *in utero* exposure to HIV infection ^55^, and in pregnant women with GBS infection and/or chorioamnionitis ^56 57^, suggesting that maternal infection can directly influence fetal CXCL8. Raised fetal inflammatory markers, therefore, do appear to be related to the maternal infection status at the time of birth, as such profiles are less obvious in infants born to mothers with recovered infection. Indeed, where we did see slight elevations in CXCL8 (and IL-6) in the recovered group, this could be explained by the mode of delivery, as labour is known to drive elevation of these cytokines ^35^.

The well-documented lymphopenia observed in COVID-19 and also in children with MIS-C ^58 59^ was not observed in infants born to mothers either with SARS-CoV-2 infection around birth or mothers who had previously had infection. This may be partially attributable to the proposed mechanisms for the observed peripheral lymphopenia, including sequestration of lymphocytes to affected organs such as the lung, direct viral invasion of T cells via ACE2 binding, or virus-induced destruction of secondary lymphoid organs ^60^ none of which apply to the infant in the absence of viral transmission. Indeed, in contrast to COVID-19 infection where reductions in peripheral NK, NKT, Vδ2 and MAIT cells have been observed in adults ^32 61 62 63^, neonates born to mothers exposed to SARS-CoV-2 actually exhibited elevated percentages of NK, NKT and Vδ2 γδ T cells and also CD161 expressing CD8 T cells (the majority of which are likely to be MAIT cells). These innate-like cells may be responding to the inflammatory cytokine milieu in the context of maternal infection potentially as a protective response in the neonate vis-à-vis their likely beneficial role in severe adult COVID-19 ^62^. Immune activation of these cells has been seen at the maternal fetal interface ^31^ and although a previous report suggested there was no elevation of NK cells in infants born to COVID-19 mothers ^29^, this was only compared to reported reference levels ^64^ and there was no direct comparator group in their study.

As well as alterations in these cell populations, we also identified enhanced cytokine potential upon *in vitro* stimulation. This was observed not only in infants born to mothers with recent/ongoing infection but also in those born to recovered mothers which suggests potential *in utero* priming of the immune response. Hence, the percentage of CD4, CD8, NK, NKT or γδ T cells that produced TNF-α (or IFN-γ and IL-17 to a lesser extent) was significantly greater in infants born to mothers exposed to SARS-CoV-2. At birth neonatal T cells predominantly produce CXCL8 with a limited capacity for IFN-γ and IL-17 production that increases with age ^65^). Thus, these findings may reflect some accelerated maturation of the neonatal immune system induced *in utero* by maternal SARS-CoV-2 infection. Much of this enhanced cytokine functionality correlated with other markers of immune maturation such as increased percentages of memory T cells and decreased T cell expression of CD38. IFN-γ expression is controlled by epigenetic mechanisms in neonates ^66^, so it is tempting to speculate that maternal SARS-CoV-2 infection may have induced some epigenetic changes in these loci. Indeed, maternal exposure to polycyclic aromatic hydrocarbons directly altered this locus in cord blood mononuclear cells ^67^ and other data suggests that epigenetic modifications during gestation can shape the future development of diseases like obesity, type 2 diabetes, allergy, asthma, and infections ^68 69^. Increased proportions of cytokine producing T cells have also been observed following *in utero* exposure to malaria, HBV and HIV-even without vertical transmission ^70-72^.

Taken together, these data strongly suggest that maternal SARS-CoV-2 infection shapes the immune profile of an infant to different extents dependent on the time of exposure. We identified a transient response to maternal inflammation in the form of increased cytokines in cord plasma but also altered immune cell functionality in neonates exposed to SARS-CoV-2 at any point during gestation, suggesting some immune imprinting. Whilst the aetiology of the observed immune perturbations in the neonate remain unclear, the consequences could be far-reaching.

More data are needed to establish if these changes specifically relate to enhanced protection from SARS-CoV-2 mediated disease or are detrimental if the infant is born in a *milieu* of inflammatory cytokines. A hyper-inflammatory MIS-C like response has been observed in a neonate following *in utero* exposure to SARS-CoV-2 (mother infected 9 weeks prior to birth) with no evidence of direct neonatal infection ^73^. Long term follow-up of the infants in our study will establish if maternal exposure to SARS-CoV-2 has a long lasting impact on the child. These data may also have implications regarding the vaccination regimen for pregnant women. Indeed, the reduced transfer of protective Abs to the infant we observed in those infants born to mothers with recent infection may suggest 2^nd^ trimester or early 3^rd^ trimester vaccination is preferable. Neonatal immune profiling following vaccination in pregnancy may also determine what level of maternal immune activation drives the neonatal imprinting observed.

## Methods

### Study design and human material

Umbilical cord blood (and paired maternal peripheral blood) was collected over the COVID-19 pandemic (28^th^ May 2020-1^st^ March 2021) at the time of birth from infants born to mothers who were SARS-CoV-2 Exposed (SE) attending the maternity unit at GSTT, London in heparinised blood tubes at the time of birth to investigate the immune status (REC Approval No. 19/SC/0232). Mother infant dyads were categorised into recent/ongoing (R/O) and recovered groups (R), according to the number of days prior to birth that the mother received a positive status from a SARS-CoV-2 nasopharyngeal swab (R/O: <14 days; R: ≥14 days). Umbilical cord blood was also collected at the time of birth from infants born to healthy mothers at GSTT prior to the COVID-19 pandemic (until 1^st^ Jan 2020), hence their mothers were not infected with SARS-CoV-2 at any time during their pregnancy, termed the Non SARS-CoV-2 Exposed (NSE) group (REC Approval No. 17/LO/0641). The clinical details of the groups can be found in Table 1.

### Isolation of CMBCs and plasma

Cord blood mononuclear cells (CBMCs) were isolated from all infant groups. Neat plasma was also isolated from the recent/ongoing and recovered groups and their paired mothers (all SE samples). More specifically, for the paired maternal peripheral blood samples, whole blood was centrifuged for 2000 *g* for 10 min at room temperature, and plasma was harvested from the upper layer and stored at -80°C in polypropylene tubes. For the neonatal samples, cord blood was layered onto Ficoll (GE Healthcare), within a 15 ml polypropylene conical tube, and centrifuged at 800 *g* for 15 min (break OFF) at room temperature. Plasma was then collected from the top fraction and stored at - 80°C in polypropylene tubes. The CBMC layer was isolated and subsequently washed twice with pre-warmed base medium (BM) (RPMI-1640 + L-Glutamine, Gibco), and then complete medium (CM) (RPMI-1640 + L-Glutamine; 10% Heat-Inactivated FBS; 1% Penicillin-Streptomycin, Gibco), under centrifugation at 300 *g* for 5 min at room temperature. The cell pellet was then frozen in Cryostor® CS10 (Sigma) within polypropylene cryovials before proceeding with flow cytometry.

### Polyclonal stimulation

Prior to staining the CBMCs for flow cytometry immunophenotyping with panel 4 (Extended Data Table 1), cells were thawed and plated in 96 well round bottom plates (Corning) within 200 μL CM containing Phorbol 12-myristate 13-acetate (PMA) (10 ng/ml) (Sigma), Ionomycin (1 μg/ml) (Sigma), Brefeldin A (20 ng/ml) (Sigma) and Monensin solution (1x) (BioLegend). A Brefeldin A and Monensin only control was also plated for each infant. Cells were incubated at 37°C for 4 h in the Biosafety level 3 (BSL-3) containment lab, in accordance with the King’s College London safety rules before proceeding with flow cytometry staining. Cells were removed from BSL-3 once they were fixed with Cell Fix (1X) (BD) for a minimum of 10 mins.

### Flow cytometry staining and acquisition

CBMCs were thawed and plated in 96 well round bottom plates before staining in one of 4 panels (Extended Data Table 1) assessing the following immune cell populations (panel 1: T cell naïve/memory status; panel 2: myeloid and B cells; panel 3: T and NK cell activation status; panel 4: T and NK cytokine potential). All four panels contained surface marker staining, and panel 1 and 4 also contained intracellular staining. All the following wash steps were performed under 2000 *g*, for 1 min at room temperature. For each panel, cells were washed with 100 μL Dulbecco’s Phosphate Buffered Saline (1X) (PBS, Gibco) and resuspended in 100 μL PBS, containing Zombie NIR™ Fixable Viability dye (1:1000) (Biolegend), with the addition of TCR Vδ1-FITC (TS8.2) (Thermo Fisher) in panel 1, for 15 mins in the dark at 4°C. Cells were then washed with 150 μL eBioscience™ FOXP3 Fixation/Permeabilization buffer (Invitrogen), for panel 1, or FACS buffer (PBS, 0.5% heat-inactivated FBS, 2mM EDTA, Invitrogen), for panels 2-4. The wash step was repeated with 200 μL volume. Cells were then resuspended in 100 μL eBioscience™ FOXP3 Permeabilization buffer (FPB) (Invitrogen), for panel 1, or 50 μL surface antibody cocktail within FACS buffer, for panels 2-4, for 30 mins (panels 2-3) or 20 mins (panel 4), at 4°C in the dark. Cells were subsequently washed in 100 μL FPB (panel 1) or 150 μL FACS buffer (panels 2-4), and again with 200 μL volume before resuspending in 50 μL antibody cocktail in FPB (panel 1), or 100 μL Cell Fix (1x) (panels 2-4) for 30 mins (panels 1-3) or until intracellular cytokine staining (ICS) (panel 4) at 4°C in the dark. Panels 1-3 were then washed twice in 100 μL FPB (panel 1) or FACS buffer (panels 2-3) and resuspended in 200 μL FACS buffer until acquisition. For the panel 4 ICS, cells were centrifuged at 2000 *g*, 1 min, at room temperature and resuspended in 50 μL ICS antibody cocktail within Permeabilization Wash Buffer (1x) (PWB) (BioLegend) for 30 mins in the dark at room temperature. Cells were washed twice in 150 μL PWB, followed by 200 μL FACS buffer before resuspending the cells in FACS buffer for acquisition in a high-throughput sampler on a 4-laser LSR Fortessa (BD), at a flow rate of 1 μL/s.

### Flow cytometry data analysis

Raw FCS files were analysed using FlowJo (v10.6.2, BD) and the gating strategies are included in Extended Data Fig. 3. As an internal control, the same adult sample was run alongside each flow cytometry experiment for consistency and to aid setting gates. Data were cleaned up by gating on the Time parameter to ensure that only cells going through a constant flow stream were analysed, and cell populations were excluded from downstream analysis if the event count in the parent population was <30.

### SARS-CoV-2 specific antibody testing

ELISAs were conducted as previously described ^30 74^. All plasma samples were heat-inactivated at 56 °C for 30 min before use. High-binding ELISA plates (Corning, 3690) were coated with antigen (Nuclear protein (N), Spike glycoprotein (S) or the receptor binding domain (RBD) at 3 µg ml^-1^ (25 µl per well) in PBS, either overnight at 4 °C or for 2 h at 37 °C. Wells were washed with PBS-T (PBS with 0.05% Tween-20) and then blocked with 100 µl of 5% milk in PBS-T for 1 h at room temperature. Wells were emptied and serial dilutions of plasma (starting at 1:25, 5-fold dilution) were added and incubated for 2 h at room temperature. Control reagents included CR3009 (2 µg/ml) (N-specific monoclonal antibody), CR3022 (0.2 µg/ml) (S-specific monoclonal antibody), negative control plasma (1:25 dilution), positive control plasma (1:50), and blank wells. Wells were washed with PBS-T. Secondary antibody was added and incubated for 1h at room temperature. IgG was detected using goat-anti-human-Fc-AP (1:1000) (Jackson, catalogue no. 109-055-098) and wells were washed with PBS-T and AP substrate (Sigma) was added and plates read at 405 nm. IgM was detected using goat-anti-human-IgM-HRP (1:1000) (Sigma catalogue no. A6907) and wells were washed with PBS-T and one-step 3,3′,5,5′-tetramethylbenzidine (TMB) substrate (Thermo Fisher Scientific) was added and quenched with 2M H_2_S0_4_ before reading at 450 nm (HRP). Samples were run in duplicate and peak fold-change above background was plotted for each patient. IgG transfer ratios were calculated to display the difference in paired infant and maternal levels [Infant IgG peak fold change over background/maternal IgG peak fold change over background].

### Quantification of plasma cytokines

Paired maternal and infant plasma were thawed and tested in the 13-plex LegendPlex Human Anti-Virus Response Panel kit (Biolegend), to quantify levels of IL-1β, IL-6, TNF-α, IP-10, CXCL8, IL-12p70, IFN-α2, IFN-λ1, IFN-λ2/3, GM-CSF, IFN-β, IL-10 and IFN-γ. The assay was performed according to the manufacturer instructions and was modified by diluting the kit reagents 2x (beads, detection antibodies and streptavidin-PE). All plasma samples were diluted 2x with assay buffer, and resulting sample concentrations were calculated according to the dilution factor. In brief, 25 μL diluted plasma or standard, and mixed beads (1:1 ratio) were added to each well (in V-bottom 96-well plates) and incubated for 1.5 h. The samples were washed twice with wash buffer, incubated with 25 μL detection antibodies for 1 h and then 25 μL streptavidin-PE was added for a further 30 mins. The samples were then washed once with wash buffer, resuspended in 200 μL wash buffer and acquired on a high-throughput sampler with a 3-laser FACSCanto™ (BD). All incubation steps were performed under 600 *rpm* on an orbital shaker at room temperature and protected from light. Data were cleaned up by excluding cytokines if the bead event count was <90, to ensure accurate analyses, performed using the Windows LegendPlex (v8.0, BioLegend) software.

### Statistical analyses

Analysed flow cytometry populations, plasma cytokines and antibodies were imported into an Excel spreadsheet and analysed in R (v4.0.3) to generate boxplots, dimensionality reduction plots, Spearman correlation plots and heatmaps. Clustered heatmaps were performed on scaled and centred data using the heatmap.2 package, and clustered according to the **Euclidean** method. The corrplot package was used to generate Spearman correlation matrices, and only significant values (p<0.05) are displayed. GraphPad Prism (v9.0) was also utilised to generate scatter plots for cytokines and the antibody heatmap. All statistical tests were used to measure the differences between biologically distinct samples. Unadjusted p values (*p<0.05; **p<0.01; ***P<0.001 and ****p<0.0001) were assessed by the Kolmogorov-Smirnov test (to compare cytokine concentrations between the groups), two-sided Wilcoxon rank-sum tests (for immune cell populations between the groups), and two-sided paired Wilcoxon tests (between paired maternal and infant antibody/cytokine levels).

## Acknowledgements

We thank the mothers and their infants for blood collection and all the midwives at STH for sample collection. We also thank Shraddha Kamdar and those staff involved with COVID-IP at King’s College London ^32^ for some sample processing and Thomas Lechmere for assistance with ELISAs. We thank Iva Zlatareva for assistance with the cytokine multiplex analysis. We would also like to acknowledge Evolve Biosystems for funding (RT and DG) towards the non SARS-CoV-2 exposed (NSE) infant cord blood sample collection and the staff involved (Niamh Kelly, Lucy McMillan, Sarah Kheirallah and Jiadai Mi). We thank Yin Wu for critical reading of the manuscript. SG is supported by a MRC-KCL Doctoral Training Partnership in Biomedical Sciences (MR/N013700/1), DLG by Action Medical Research (GN2790), MC is supported by the NIHR BRC COVID-19 call, RMT by Tommy’s (Charity No. 1060508) and Borne (1167073) and JS and KJD by Kings Together Rapid COVID-19 call award and Huo Family Foundation award. The research was also supported by the National Institute for Health Research (NIHR) Biomedical Research Centre based at Guy’s and St Thomas’ NHS Foundation Trust (GSTT) and King’s College London (KCL) (part of the King’s Health Partners Academic Sciences Centre). The views expressed are those of the authors and not necessarily those of the NHS, the NIHR or the Department of Health.

## Author Contributions

MC and CM: Patient consent, study design and sample collection. SG: Sample processing, flow cytometry and data analysis. SG and AD: panel design, multiplex cytokine testing and analysis. JS and KJD: Antibody ELISA testing and analysis. DG and RT: Manuscript writing and experimental design. All authors reviewed drafts of the manuscript prior to submission.

## Competing Interests

The authors declare no competing interests.

## Notes

### Competing Interest Statement

The authors have declared no competing interest.

## References

1. Sutton, D., Fuchs, K., D’Alton, M. & Goffman, D. Universal Screening for SARS-CoV-2 in Women Admitted for Delivery. N Engl J Med (2020).

2. Chen, L., et al. Clinical Characteristics of Pregnant Women with Covid-19 in Wuhan, China. N Engl J Med 382, e100 (2020).

3. Pierce-Williams, R.A.M., et al. Clinical course of severe and critical coronavirus disease 2019 in hospitalized pregnancies: a United States cohort study. Am J Obstet Gynecol MFM 2, 100134 (2020).

4. Lokken, E.M., et al. Clinical characteristics of 46 pregnant women with a severe acute respiratory syndrome coronavirus 2 infection in Washington State. Am J Obstet Gynecol 223, 911 e911–911 e914 (2020).

5. Allotey, J., et al. Clinical manifestations, risk factors, and maternal and perinatal outcomes of coronavirus disease 2019 in pregnancy: living systematic review and meta-analysis. BMJ 370, m3320 (2020).

6. Kadiwar, S., et al. Were pregnant women more affected by COVID-19 in the second wave of the pandemic? Lancet 397, 1539–1540 (2021).

7. Vousden, N., et al. The incidence, characteristics and outcomes of pregnant women hospitalized with symptomatic and asymptomatic SARS-CoV-2 infection in the UK from March to September 2020: A national cohort study using the UK Obstetric Surveillance System (UKOSS). PloS one 16, e0251123 (2021).

8. Martinez-Perez, O., et al. The association between SARS-CoV-2 infection and preterm delivery: a prospective study with a multivariable analysis. BMC Pregnancy Childbirth 21, 273 (2021).

9. Gale, C., et al. Characteristics and outcomes of neonatal SARS-CoV-2 infection in the UK: a prospective national cohort study using active surveillance. Lancet Child Adolesc Health 5, 113–121 (2021).

10. Salvatore, C.M., et al. Neonatal management and outcomes during the COVID-19 pandemic: an observation cohort study. Lancet Child Adolesc Health 4, 721–727 (2020).

11. Diriba, K., Awulachew, E. & Getu, E. The effect of coronavirus infection (SARS-CoV-2, MERS-CoV, and SARS-CoV) during pregnancy and the possibility of vertical maternal-fetal transmission: a systematic review and meta-analysis. Eur J Med Res 25, 39 (2020).

12. Vivanti, A.J., et al. Transplacental transmission of SARS-CoV-2 infection. Nat Commun 11, 3572 (2020).

13. Zeng, H., et al. Antibodies in Infants Born to Mothers With COVID-19 Pneumonia. JAMA (2020).

14. Lorenz, N., et al. Neonatal Early-Onset Infection With SARS-CoV-2 in a Newborn Presenting With Encephalitic Symptoms. Pediatr Infect Dis J 39, e212 (2020).

15. Fenizia, C., et al. Analysis of SARS-CoV-2 vertical transmission during pregnancy. Nat Commun 11, 5128 (2020).

16. Shende, P., et al. Persistence of SARS-CoV-2 in the first trimester placenta leading to transplacental transmission and fetal demise from an asymptomatic mother. Hum Reprod 36, 899–906 (2021).

17. Dong, L., et al. Possible Vertical Transmission of SARS-CoV-2 From an Infected Mother to Her Newborn. JAMA 323, 1846–1848 (2020).

18. Raschetti, R., et al. Synthesis and systematic review of reported neonatal SARS-CoV-2 infections. Nat Commun 11, 5164 (2020).

19. Chen, G., et al. Immune Response to COVID-19 During Pregnancy. Front Immunol 12, 675476 (2021).

20. Garcia-Flores, V., et al. Maternal-Fetal Immune Responses in Pregnant Women Infected with SARS-CoV-2. Res Sq (2021).

21. Kamdar, S., et al. Perinatal inflammation influences but does not arrest rapid immune development in preterm babies. Nat Commun 11, 1284 (2020).

22. Hygino, J., et al. Enhanced Th17 phenotype in uninfected neonates born from viremic HIV-1-infected pregnant women. J Clin Immunol 31, 186–194 (2011).

23. Gabriel, B., et al. Analysis of the TCR Repertoire in HIV-Exposed but Uninfected Infants. Sci Rep 9, 11954 (2019).

24. Abu-Raya, B., Kollmann, T.R., Marchant, A. & MacGillivray, D.M. The Immune System of HIV-Exposed Uninfected Infants. Front Immunol 7, 383 (2016).

25. Babik, J.M., Cohan, D., Monto, A., Hartigan-O’Connor, D.J. & McCune, J.M. The human fetal immune response to hepatitis C virus exposure in utero. J Infect Dis 203, 196–206 (2011).

26. Gomez de Aguero, M., et al. The maternal microbiota drives early postnatal innate immune development. Science 351, 1296–1302 (2016).

27. Arrieta, M.C., et al. Early infancy microbial and metabolic alterations affect risk of childhood asthma. Sci Transl Med 7, 307ra152 (2015).

28. Torow, N. & Hornef, M.W. The Neonatal Window of Opportunity: Setting the Stage for Life-Long Host-Microbial Interaction and Immune Homeostasis. J Immunol 198, 557–563 (2017).

29. Liu, P., et al. The immunologic status of newborns born to SARS-CoV-2-infected mothers in Wuhan, China. J Allergy Clin Immunol 146, 101–109 e101 (2020).

30. Pickering, S., et al. Comparative assessment of multiple COVID-19 serological technologies supports continued evaluation of point-of-care lateral flow assays in hospital and community healthcare settings. PLoS Pathog 16, e1008817 (2020).

31. Lu-Culligan, A., et al. Maternal respiratory SARS-CoV-2 infection in pregnancy is associated with a robust inflammatory response at the maternal-fetal interface. Med (N Y) 2, 591–610 e510 (2021).

32. Laing, A.G., et al. A dynamic COVID-19 immune signature includes associations with poor prognosis. Nature medicine 26, 1623–1635 (2020).

33. Lucas, C., et al. Longitudinal analyses reveal immunological misfiring in severe COVID-19. Nature 584, 463–469 (2020).

34. Arunachalam, P.S., et al. Systems biological assessment of immunity to mild versus severe COVID-19 infection in humans. Science 369, 1210–1220 (2020).

35. Tornblom, S.A., et al. mRNA expression and localization of bNOS, eNOS and iNOS in human cervix at preterm and term labour. Reprod Biol Endocrinol 3, 33 (2005).

36. Gleditsch, D.D., et al. Maternal inflammation modulates infant immune response patterns to viral lung challenge in a murine model. Pediatric research 76, 33–40 (2014).

37. Lacorcia, M. & Prazeres da Costa, C.U. Maternal Schistosomiasis: Immunomodulatory Effects With Lasting Impact on Allergy and Vaccine Responses. Front Immunol 9, 2960 (2018).

38. Straubinger, K., et al. Maternal immune response to helminth infection during pregnancy determines offspring susceptibility to allergic airway inflammation. J Allergy Clin Immunol 134, 1271–1279 e1210 (2014).

39. Apostol, A.C., Jensen, K.D.C. & Beaudin, A.E. Training the Fetal Immune System Through Maternal Inflammation-A Layered Hygiene Hypothesis. Front Immunol 11, 123 (2020).

40. Bilbo, S.D. & Schwarz, J.M. Early-life programming of later-life brain and behavior: a critical role for the immune system. Front Behav Neurosci 3, 14 (2009).

41. Williamson, L.L., et al. Got worms? Perinatal exposure to helminths prevents persistent immune sensitization and cognitive dysfunction induced by early-life infection. Brain Behav Immun 51, 14–28 (2016).

42. Helgertz, J. & Bengtsson, T. The Long-Lasting Influenza: The Impact of Fetal Stress During the 1918 Influenza Pandemic on Socioeconomic Attainment and Health in Sweden, 1968-2012. Demography 56, 1389–1425 (2019).

43. Acquah, J.K., Dahal, R. & Sloan, F.A. 1918 Influenza Pandemic: In Utero Exposure in the United States and Long-Term Impact on Hospitalizations. Am J Public Health 107, 1477–1483 (2017).

44. Flannery, D.D., et al. Assessment of Maternal and Neonatal Cord Blood SARS-CoV-2 Antibodies and Placental Transfer Ratios. JAMA Pediatr (2021).

45. Munoz, F.M., et al. Safety and immunogenicity of tetanus diphtheria and acellular pertussis (Tdap) immunization during pregnancy in mothers and infants: a randomized clinical trial. JAMA 311, 1760–1769 (2014).

46. Martinez, D.R., et al. Fc Characteristics Mediate Selective Placental Transfer of IgG in HIV-Infected Women. Cell 178, 190–201 e111 (2019).

47. Fouda, G.G., Martinez, D.R., Swamy, G.K. & Permar, S.R. The Impact of IgG transplacental transfer on early life immunity. Immunohorizons 2, 14–25 (2018).

48. Goncalves, G., et al. Transplacental transfer of measles and total IgG. Epidemiol Infect 122, 273–279 (1999).

49. Atyeo, C., et al. Compromised SARS-CoV-2-specific placental antibody transfer. Cell 184, 628–642 e610 (2021).

50. Bordt, E.A., et al. Sexually dimorphic placental responses to maternal SARS-CoV-2 infection. bioRxiv (2021).

51. Beharier, O., et al. Efficient Maternal to Neonatal transfer of SARS-CoV-2 and BNT162b2 antibodies. MedRxiv (2021).

52. Rottenstreich, A., et al. Efficient maternofetal transplacental transfer of anti-SARS-CoV-2 spike antibodies after antenatal SARS-CoV-2 BNT162b2 mRNA vaccination. Clin Infect Dis (2021).

53. Lu, W., et al. Early immune responses and prognostic factors in children with COVID-19: a single-center retrospective analysis. BMC Pediatr 21, 181 (2021).

54. Sherer, M.L., et al. Dysregulated immunity in SARS-CoV-2 infected pregnant women. medRxiv (2020).

55. Lohman-Payne, B., et al. HIV-exposed uninfected infants: elevated cord blood Interleukin 8 (IL-8) is significantly associated with maternal HIV infection and systemic IL-8 in a Kenyan cohort. Clin Transl Med 7, 26 (2018).

56. Ren, J., Qiang, Z., Li, Y.Y. & Zhang, J.N. Biomarkers for a histological chorioamnionitis diagnosis in pregnant women with or without group B streptococcus infection: a case-control study. BMC Pregnancy Childbirth 21, 250 (2021).

57. Reuschel, E., et al. Perinatal Gram-Positive Bacteria Exposure Elicits Distinct Cytokine Responses In Vitro. Int J Mol Sci 22(2020).

58. Consiglio, C.R., et al. The Immunology of Multisystem Inflammatory Syndrome in Children with COVID-19. Cell 183, 968–981 e967 (2020).

59. Carter, M.J., et al. Peripheral immunophenotypes in children with multisystem inflammatory syndrome associated with SARS-CoV-2 infection. Nature medicine 26, 1701–1707 (2020).

60. Yang, L., et al. COVID-19: immunopathogenesis and Immunotherapeutics. Signal Transduct Target Ther 5, 128 (2020).

61. Kuri-Cervantes, L., et al. Comprehensive mapping of immune perturbations associated with severe COVID-19. Sci Immunol 5(2020).

62. Jouan, Y., et al. Phenotypical and functional alteration of unconventional T cells in severe COVID-19 patients. J Exp Med 217(2020).

63. Sun, D., et al. SARS-CoV-2 infection in infants under 1 year of age in Wuhan City, China. World J Pediatr 16, 260–266 (2020).

64. Amatuni, G.S., et al. Reference intervals for lymphocyte subsets in preterm and term neonates without immune defects. J Allergy Clin Immunol 144, 1674–1683 (2019).

65. Gibbons, D., et al. Interleukin-8 (CXCL8) production is a signatory T cell effector function of human newborn infants. Nature medicine 20, 1206–1210 (2014).

66. White, G.P., Watt, P.M., Holt, B.J. & Holt, P.G. Differential patterns of methylation of the IFN-gamma promoter at CpG and non-CpG sites underlie differences in IFN-gamma gene expression between human neonatal and adult CD45RO-T cells. J Immunol 168, 2820–2827 (2002).

67. Tang, W.Y., et al. Maternal exposure to polycyclic aromatic hydrocarbons and 5’-CpG methylation of interferon-gamma in cord white blood cells. Environ Health Perspect 120, 1195–1200 (2012).

68. Sureshchandra, S., et al. Phenotypic and Epigenetic Adaptations of Cord Blood CD4+ T Cells to Maternal Obesity. Front Immunol 12, 617592 (2021).

69. van Esch, B., et al. The Impact of Milk and Its Components on Epigenetic Programming of Immune Function in Early Life and Beyond: Implications for Allergy and Asthma. Front Immunol 11, 2141 (2020).

70. Odorizzi, P.M., et al. In utero priming of highly functional effector T cell responses to human malaria. Sci Transl Med 10(2018).

71. Hong, M., et al. Trained immunity in newborn infants of HBV-infected mothers. Nat Commun 6, 6588 (2015).

72. Garcia-Knight, M.A., et al. Altered Memory T-Cell Responses to Bacillus Calmette-Guerin and Tetanus Toxoid Vaccination and Altered Cytokine Responses to Polyclonal Stimulation in HIV-Exposed Uninfected Kenyan Infants. PloS one 10, e0143043 (2015).

73. Kappanayil, M., et al. Multisystem inflammatory syndrome in a neonate, temporally associated with prenatal exposure to SARS-CoV-2: a case report. Lancet Child Adolesc Health 5, 304–308 (2021).

74. Seow, J., et al. Longitudinal observation and decline of neutralizing antibody responses in the three months following SARS-CoV-2 infection in humans. Nat Microbiol 5, 1598–1607 (2020).

