## Extended Data Figs and Table for "The legacy of maternal SARS-CoV-2 infection on the immunology of the neonate"

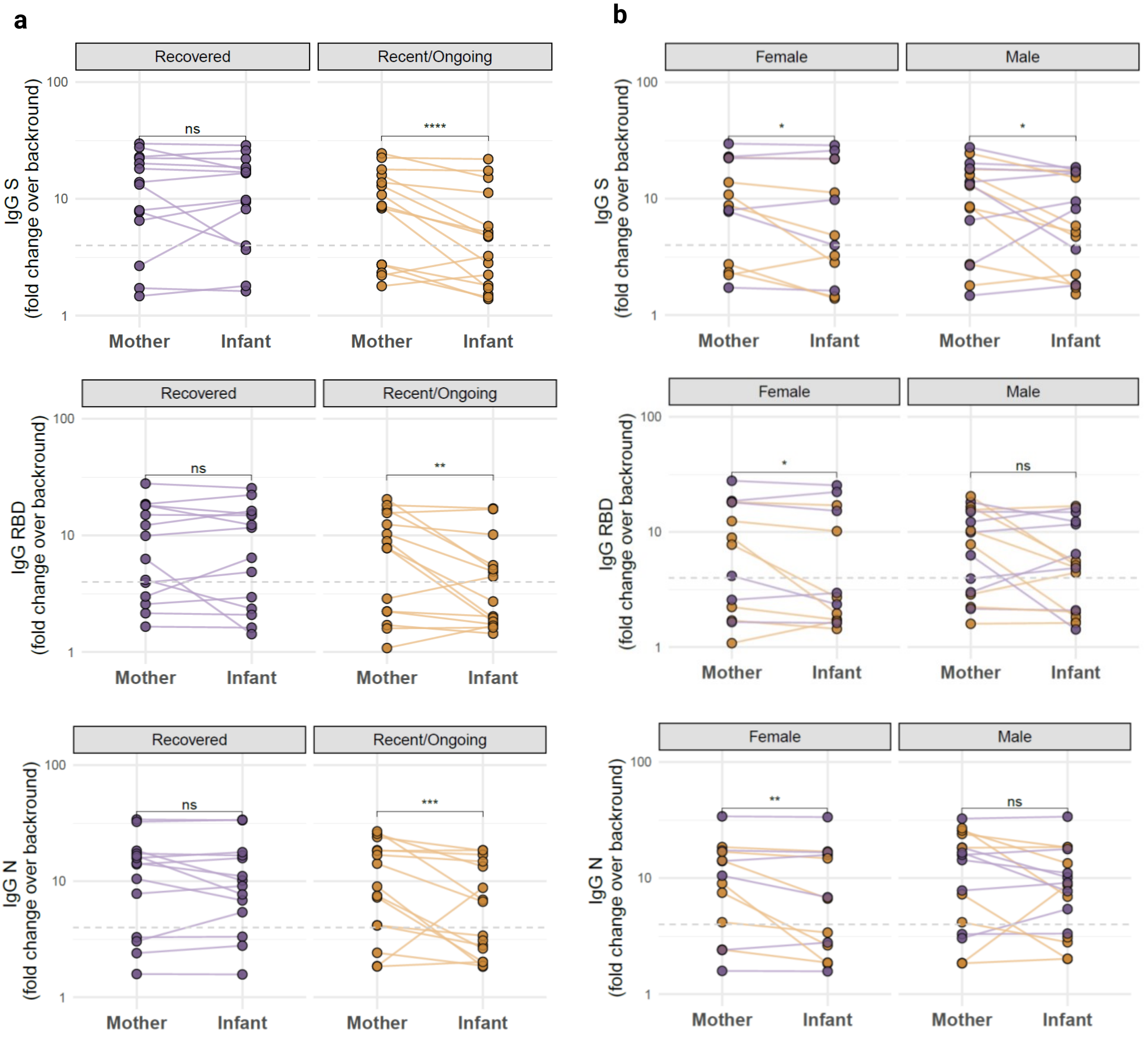


**Extended Data Fig. 1 | Infant IgG levels to all three SARS-CoV-2 epitopes were lower than their mothers in the recent/ongoing group and a sex bias was not observed. a-b**, Infant and mother paired plasma IgG levels against the S, RBD and N epitopes across the R (*n=14*) and R/O (*n=15*) groups (**a**), and in female (*n=13*) and male (*n=16*) infants (**b**). Peak IgG/IgM levels (fold change over background) are plotted within the infant-mother dyads. Seropositive IgG threshold (dashed grey line) = 4x fold change over background. Each line joins an infant and their paired mother. *p<0.05; **p<0.01; ***<p<0.001; ****p<0.0001 were assessed by two-sided paired Wilcoxon tests.

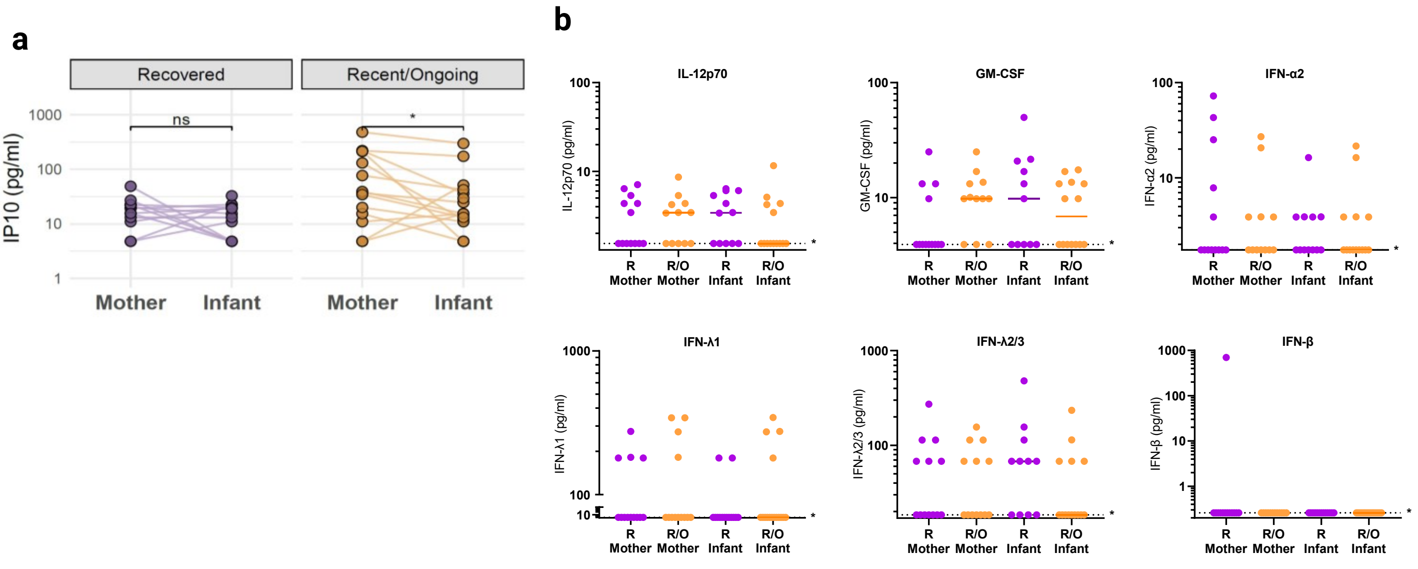


**Extended Data Fig. 2 | Plasma cytokine concentrations in infants and their mothers within the recent/ongoing and recovered groups. a**, Paired maternal and infant plasma IP-10 levels within the R (*n=12*) and the R/O (*n=14*) groups. Each line joins an infant and their paired mother. **b**, Plasma IL-12p70, GM-CSF, IFN-α2, IFN-λ1, IFN-λ2/3 and IFN-β in the mothers (R - *n=13*, R/O - IFN-λ1, IFN-λ2/3: *n=13*, GM-CSF, IL-12p70, IFN-β: *n=12*) and their infants (R - IFN-λ1: *n=14*, IFN-β: *n=13*, IL-12p70, IFN-α2: *n=12*, GM-CSF: *n=11*, R/O - IFN-λ1: *n=16*, IFN-β: *n=15*, IFN-λ2/3, GM-CSF, IL-12p70: *n=14*). TNFα and IFNγ cytokines were not detected in any samples and are not presented. Horizontal dotted lines represent the minimum detectable concentrations. P values (*p<0.05) were assessed by two-sided paired Wilcoxon tests.

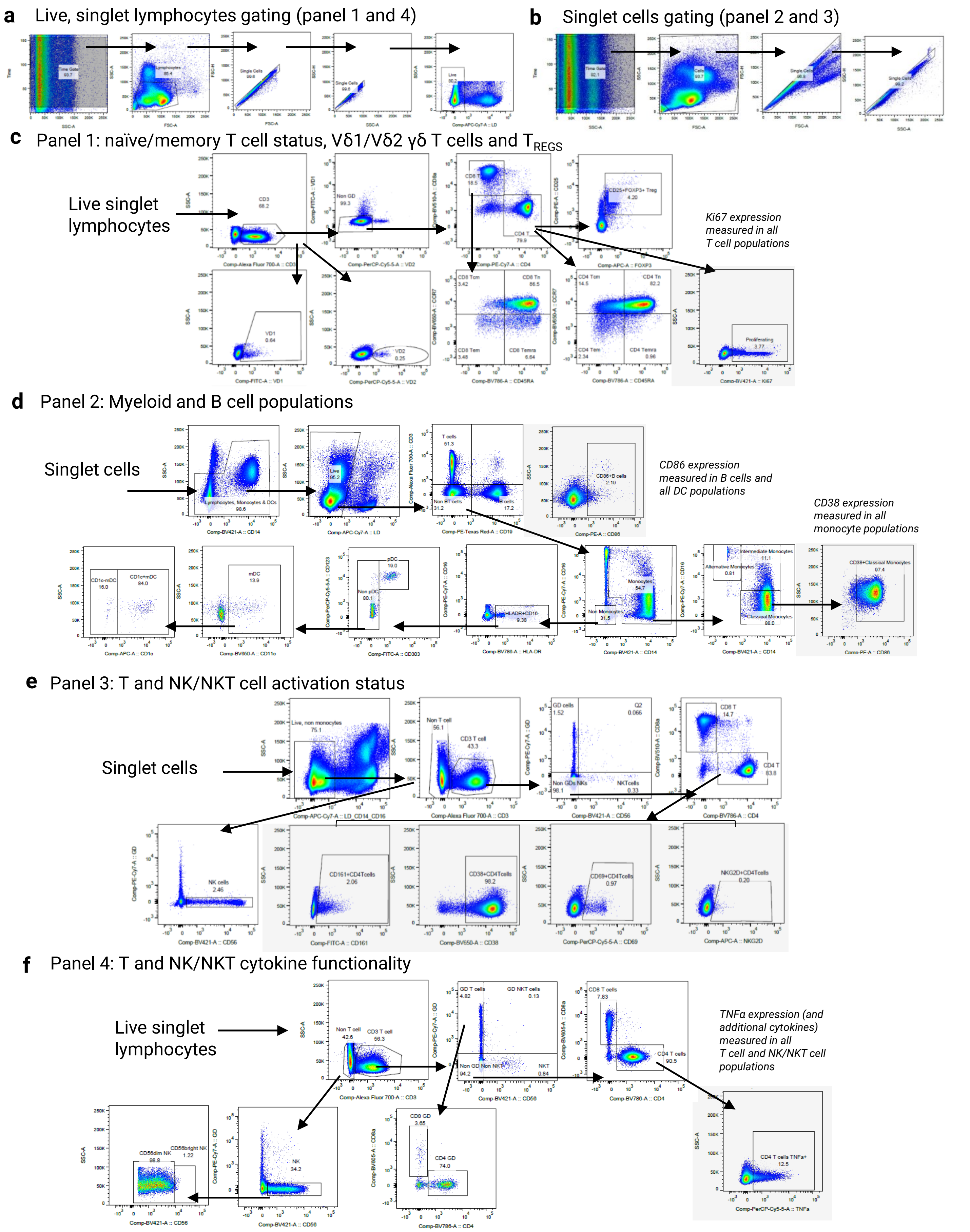


**Extended Data Fig. 3 | Flow cytometry gating strategies for all four panels. a-b**, Representative initial gating strategies for all 4 panels. Live singlet lymphocytes (**a**) or cells (**b**) were gated on Time vs SSC-A to ensure that only cells going through a constant flow stream were analysed. **c-f**, Representative gating strategies for panels 1-4 after initial gating. T cell naïve/memory status, Vδ1/Vδ2 γδ T cell subtypes and T$_{REGS}$ (and their Ki67 expression) (**c**), B cell, dendritic cell (DC) an monocyte populations (and their CD86 or CD38 expression) (**d**), T cell and NK/NKT cell activation status (**e**) and T cell/NK/NKT cytokine functionality after polyclonal stimulation (**f**) were measured. Plots in grey boxes indicate populations that were measured in multiple cell types as described.

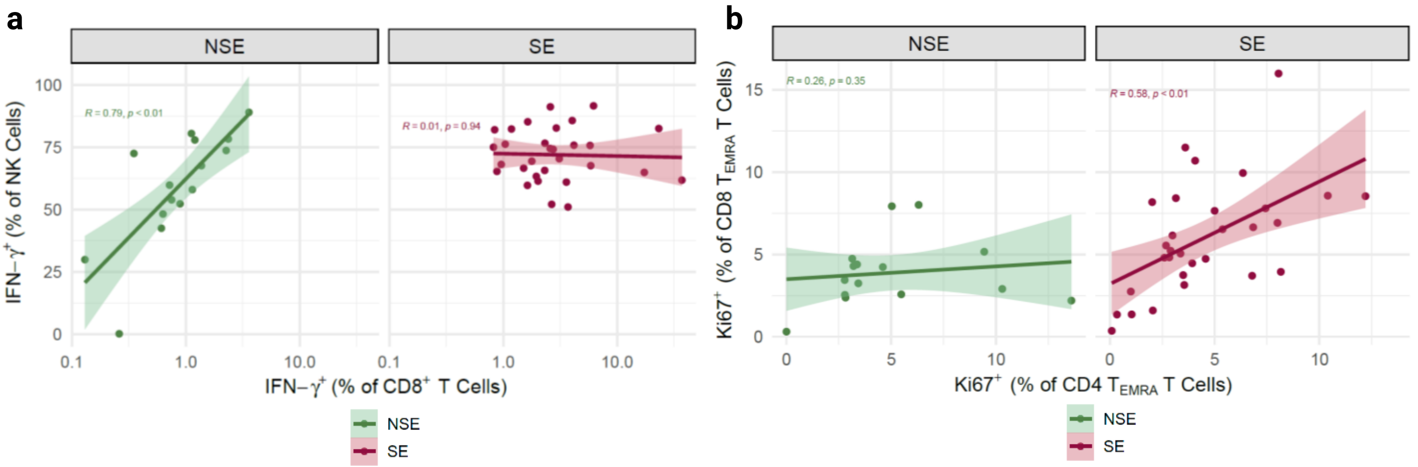


**Extended Data Fig. 4 | Disrupted infant immune cell correlations within the SARS-CoV-2 exposed cohort. a-b**, Spearman correlation plots between Ki67+CD8+ $T_{EMRA}$ and Ki67+CD8+ $T_{EMRA}$ populations (**a**) and IFNγ+CD8+ T cells and IFNγ+ NK cells (**b**) in the NSE (*n=15*) and SE (*n=30*) infant cohorts with generalised linear model lines and 95% confidence intervals.

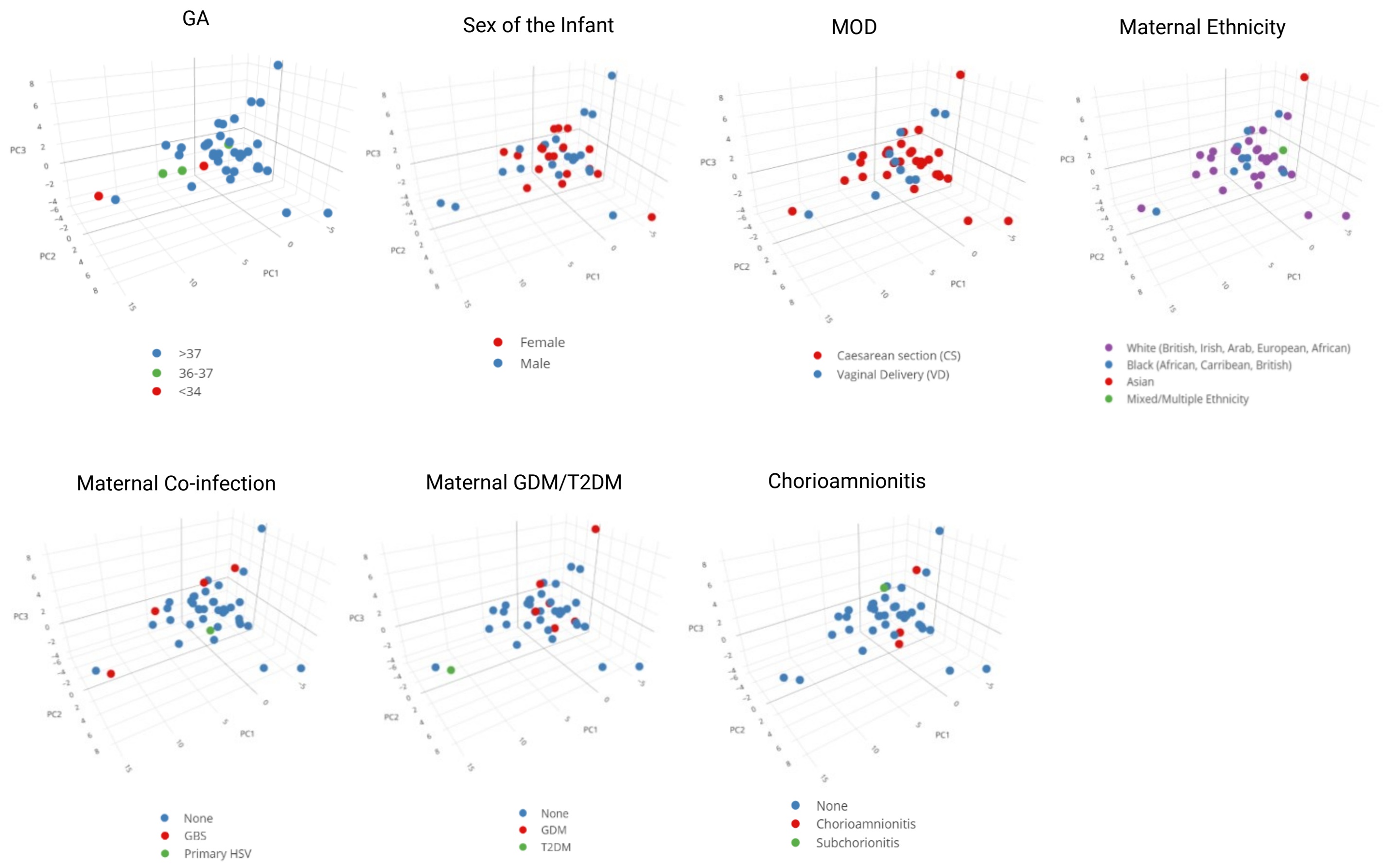


**Extended Data Fig. 5 | PCA of the flow cytometry analysis coloured by alternative factors.** 3-dimensional PCA dimensionality reduction of all flow cytometry data in the NSE (*n=13*) and SE (*n=28*) infant cohorts, coloured by alternative factors (GA, sex of the infant, MOD, maternal ethnicity, maternal co-infection, maternal GDM/T2DM and chorioamnionitis). Chorioamnionitis was defined by placental histology testing: Individuals in the 'none' group were either not tested (as no placental pathology suspected) or normal placental histology upon testing. PC1, PC2, and PC3 explain 20.6%, 11.8% and 8.4% of the variance, respectively.

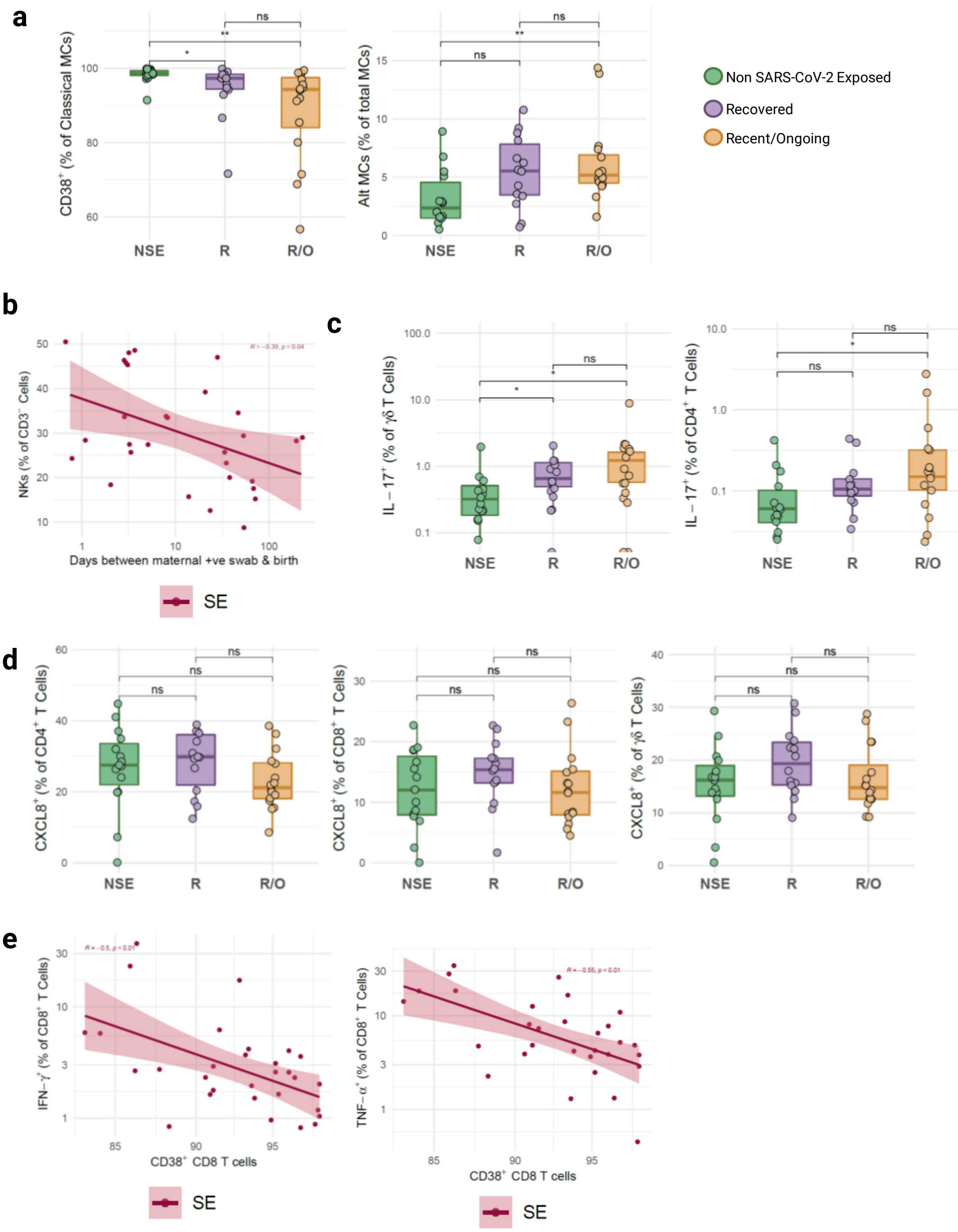


**Extended Data Fig. 6| Altered and unchanged cell populations in infants within the SARS-CoV-2 exposed cohort. a**, Boxplots displaying the proportions of $CD38^+$ classical monocytes and alternative monocytes in infants within NSE (*n=14*), R (*n=14*) and R/O (*n=16*) groups. **b**, Spearman correlation plots in the SE cohort (*n=30*) of infant proportions of NK cells and days between maternal positive nasopharyngeal swab and birth with generalised linear model lines and 95% confidence intervals. **c**, Boxplots displaying the proportions of IL-$17^+$ γδ T cells and $CD4^+$ αβ T cells in infants within NSE (*n=15*), R (*n=14*) and R/O (*n=16*) groups. **d**, Boxplots displaying the proportions f $CXCL8^+$ $CD4^+$ αβ T cells, $CD8^+$ αβ T cells and γδ T cells in infants within NSE (*n=15*), R (*n=14*) and R/O (*n=16*) groups. **e**, Spearman correlation plots in the SE cohort (*n=30*) of infant proportions of IFN-$γ^+$ (LHS) and TNF-$α^+$ (RHS) $CD8^+$ T cells and $CD38^+CD8^+$ T cells with generalised linear model lines and 95% confidence intervals. All boxplots follow standard Tukey representations; central line = median, upper line = $75^{th}$ percentile; lower line = $25^{th}$ line; whiskers = $1.5*75^{th}/25^{th}$ percentile. Unadjusted p values (*$p<0.05$ and **$p<0.01$) were assessed by two-sided Wilcoxon rank-sum tests.

| Panel 1 | | | | | |
|---|---|---|---|---|---|
| **Marker** | Fluorophore | Supplier | Cat No. | Clone | Dilution |
| **CD45RA** | BV785 | BioLegend | 304140 | HI100 | 1/100 |
| **CCR7** | BV650 | BioLegend | 353234 | GO43H7 | 1/50 |
| **CD8a** | BV510 | BioLegend | 301048 | RPA-T8 | 1/400 |
| **Ki-67** | BV421 | BioLegend | 350506 | Ki-67 | 1/50 |
| **CD4** | PE-Cy7 | BioLegend | 317414 | OKT4 | 1/50 |
| **CD25** | PE | BioLegend | 356104 | M-A251 | 1/50 |
| **TCR Vδ2** | PerCP | BioLegend | 331410 | B6 | 1/50 |
| **TCR Vδ1** | FITC | Thermo Scientific | TCR2730 | TS8.2 | 1/100 |
| **FOXP3** | AF647 | BioLegend | 320214 | 259D | 1/50 |
| **CD3** | AF700 | BioLegend | 317340 | OKT3 | 1/200 |
| **Human TruStain FcX™** | N/A | BioLegend | 422302 | N/A | 2.5/50 |
| **Panel 2** | | | | | |
| **Marker** | Fluorophore | Supplier | Cat No. | Clone | Dilution |
| **HLA-DR** | BV785 | BioLegend | 307642 | L243 | 1/100 |
| **CD11c** | BV650 | BioLegend | 301638 | 3.9 | 1/50 |
| **CD40** | BV510 | BioLegend | 334330 | 5C3 | 1/100 |
| **CD14** | BV421 | BioLegend | 301830 | M5E2 | 1/100 |
| **CD16** | PE-Cy7 | BioLegend | 302016 | 3G8 | 1/200 |
| **CD19** | PE-Dazzle 594 | BioLegend | 302252 | HIB19 | 1/100 |
| **CD86** | PE | BioLegend | 305406 | IT2.2 | 1/100 |
| **CD123** | PerCP-Cy5.5 | BioLegend | 306016 | 6H6 | 1/50 |
| **CD303** | FITC | BioLegend | 348208 | 201A | 1/100 |
| **CD1c (BDCA-1)** | APC | BioLegend | 331524 | L161 | 1/100 |
| **CD3** | AF700 | BioLegend | 317340 | OKT3 | 1/200 |
| **Human TruStain FcX™** | N/A | BioLegend | 422302 | N/A | 2.5/50 |
| **Panel 3** | | | | | |
| **Marker** | Fluorophore | Supplier | Cat No. | Clone | Dilution |
| **CD4** | BV785 | BioLegend | 317442 | OKT4 | 1/50 |
| **CD38** | BV650 | BioLegend | 356620 | HB-7 | 1/100 |
| **CD8a** | BV510 | BioLegend | 301048 | RPA-T8 | 1/400 |
| **CD56** | BV421 | BioLegend | 362552 | 5.1H11 | 1/200 |
| **TCR γδ** | PE-Cy7 | BioLegend | 331222 | B1 | 1/50 |
| **CD69** | PerCP-Cy5.5 | BioLegend | 310926 | FN50 | 1/50 |
| **CD161** | AF488 | BioLegend | 339924 | HP-3G10 | 1/50 |
| **NKG2D** | APC | BioLegend | 320808 | 1D11 | 1/50 |
| **CD3** | AF700 | BioLegend | 317340 | OKT3 | 1/200 |
| **CD16** | APC-Cy7 | BioLegend | 302018 | 3G8 | 1/100 |
| **CD14** | APC-Cy7 | BioLegend | 325620 | HCD14 | 1/100 |
| **Human TruStain FcX™** | N/A | BioLegend | 422302 | N/A | 2.5/50 |
| **Panel 4** | | | | | |
| **Marker** | Fluorophore | Supplier | Cat No. | Clone | Dilution |
| **CD4** | BV785 | BioLegend | 317442 | OKT4 | 1/50 |
| **IFN-γ** | BV650 | BioLegend | 502538 | 4S.B3 | 1/50 |
| **CD8a** | BV605 | BioLegend | 301040 | RPA-T8 | 1/100 |
| **IL-2** | BV510 | BioLegend | 500338 | MQ1-17H12 | 1/50 |
| **CD56** | BV421 | BioLegend | 362552 | 5.1H11 | 1/200 |
| **TCR γδ** | PE-Cy7 | BioLegend | 331222 | B1 | 1/50 |
| **IL-10** | PE-Dazzle 594 | BioLegend | 501426 | JES3-9D7 | 1/50 |
| **IL-4** | PE | BioLegend | 500705 | 8D4-8 | 1/50 |
| **TNF-α** | PerCP-Cy5.5 | BioLegend | 502926 | MAb11 | 1/50 |
| **IL-8** | FITC | BioLegend | 511406 | E8N1 | 1/50 |
| **CD3** | AF700 | BioLegend | 317340 | OKT3 | 1/200 |
| **IL-17A** | AF647 | BioLegend | 512310 | BL168 | 1/50 |
| **IL-17F** | AF647 | BD | 561333 | 033-782 | 1/50 |

**Extended Data Table. 1 | Flow cytometry antibodies.**
